# Cross-Species Insights from ART-D to Uncover Evolutionarily Conserved Oncogenic Mechanisms

**DOI:** 10.1101/2025.10.20.683390

**Authors:** Yifan Guo, Jiadong Zheng, Yixin Wu, Kun Zhao, Dian Lv, Wenhan Liu, Mengling Wang, Jin-Yu Lu, Wenyan Xu, Xianping Wang, Xianjue Ma

## Abstract

Cancer arises from oncogenic clones, yet the dynamic mechanisms governing their stepwise evolution toward malignancy remain incompletely understood. Here, we establish the Atlas of *Ras*-driven Tumors in *Drosophila* (ART-D), a systematic, cross-species platform that dissects the molecular and phenotypic trajectories of tumorigenesis through ten genetically defined *Ras^V12^*-driven models. By integrating longitudinal phenotypic profiling, we define three conserved stages of tumor development—initiation, promotion, and progression—distinguished by dynamic changes in tumor burden and tumor-induced cachexia. Transcriptomic dynamics reveal stage-specific signaling rewiring: early tumorigenesis is marked by co-activation of JAK/STAT, NF-κB/Toll, and MAPK pathways, whereas malignant progression is driven by Notch hyperactivation and Hippo pathway inactivation. Through integrative multi-omics and machine learning, we uncover an evolutionarily conserved pathogenic network involving coordinated JNK, NF-κB/Toll, Notch, and Hippo signaling that is functionally validated across species. ART-D provides a transformative resource that bridges *Drosophila* genetics and human cancer biology, offering a framework for decoding conserved oncogenic principles and enabling precision targeting of stage-specific vulnerabilities in *RAS*-driven cancers.

## Introduction

Cancer initiation is driven by oncogenic mutations that propel clonal evolution ^1–4^, with *RAS* proto-oncogenes representing the most frequently mutated oncogene family across human malignancies ^5–8^. Recurrent mutations at codons 12, 13, and 61 lock RAS in a constitutively active state, leading to persistent downstream signaling and uncontrolled cellular proliferation ^5–8^. Despite advances in mapping cancer genomes and identifying co-occurring or mutually exclusive mutations (e.g., *RAS-BRAF* exclusivity or *RAS-MET*/*ERBB2*-amplifications co-occurrence) ^9–13^, our understanding of how RAS dynamically interfaces with tumor suppressor networks to orchestrate malignant transformation remains fundamentally limited. Current *in vitro* models often fail to recapitulate the spatiotemporal complexity of tumor evolution *in vivo*, particularly the dynamic transitions underlying clinical progression

In this context, *Drosophila melanogaster* emerges as a uniquely powerful system for dissecting tumorigenesis. Compared to mammalian models such as *Mus musculus* or *Danio rerio*, the fruit fly offers unparalleled advantages in genetic tractability, rapid generation time, and scalability for high-throughput functional screening. Over the past decades, *Drosophila* research has been instrumental in defining foundational principles of cancer biology, from the discovery of hyperplastic/neoplastic tumor suppressors ^14^, including core members of Hippo pathway (*hippo* [*hpo*] and *warts* [*wts*]) ^15–21^ and cell polarity genes (*scribble* [*scrib*], *discs large 1* [*dlg1*], and *lethal (2) giant larvae* [*l(2)gl*]) ^22–26^, to the identification of pro-tumoral signals like c-Jun N-terminal kinase (JNK) and Notch ^14, 27–32^. Combining oncogenic *Ras* (*Ras^V12^*) with specific cooperating mutations enables the faithful modeling of hallmark cancer phenotypes, including uncontrolled proliferation, tissue invasion, systemic cachexia, and therapy resistance ^32–34^. For instance, co-expression of *Ras^V12^* with dysregulated *fiery mountain* (*fmt*), *emei* (*emei*), or *misshapen* (*msn*) activate JNK while inactivate Hippo to promote tumorigenesis and invasiveness ^35–38^. Meanwhile, *Ras^V12^* tumors with disrupted apical-basal polarity (e.g., *scrib* and *dlg*) secrete cachectic factors like Matrix metalloproteinase 1 (MMP1) and Ecdysone-inducible gene L2 (ImpL2) to remodel microenvironments and induce organism-wide wasting ^39–41^. Despite these advances, a systematic, comparative framework for *RAS*-driven tumor models with diverse genetic backgrounds has been lacking. This gap has obscured the identification of conserved molecular trajectories and stage-specific vulnerabilities that could inform targeted therapeutic strategies.

A central challenge in cancer biology lies in decoding the molecular and phenotypic transitions that define tumor evolution—from initial clonal expansion to heterogeneous, invasive disease. This complexity hinders the translation of genotype into clinically meaningful outcomes ^4^. Longitudinal analyses that integrate transcriptional dynamics with quantitative phenotypic readouts—such as tumor burden and systemic cachexia—are rare but essential for pinpointing critical intervention windows. To address this, we developed the Atlas of *Ras-*driven Tumors in *Drosophila* (ART-D), a cross-species platform encompassing ten genetically defined malignant models. By integrating deep phenotyping with longitudinal transcriptomics, we chart the conserved signaling trajectories that underlie tumorigenesis. Strikingly, multi-omics and machine learning analyses reveal that *Drosophila Ras*-driven tumors converge on a pathogenic network involving coordinated dysregulation of JNK, Notch,

NF-κB/Toll, and Hippo pathways—a signature robustly mirrored in *KRAS*/*NRAS*-high human cancers. Our work establishes *Drosophila* as a dynamic, scalable model for dissecting the evolutionary trajectory of RAS-driven tumors and uncovers evolutionarily ancient, therapeutically targetable nodes in cancer progression.

## Results

### Tumor Suppressors Synergizing with *Ras^V12^* to Drive Tumorigenesis

We previously conducted a large-scale ethyl methanesulfonate (EMS)-induced forward genetic screen on *Drosophila* chromosome 3L using *ey*-FLP-based MACRM (mosaic analysis with repressible cell marker) system (Fig. S1A) ^35, 42^, with the aim of identifying novel tumor suppressors capable of synergizing with oncogenic *Ras* (*Ras^V12^*) to drive malignant tumors in the *Drosophila* eye-antennal disc ^35, 36, 38, 42^. Among the candidate alleles identified, eight recessive-lethal alleles were characterized: *Vacuolar protein sorting 36* (*Vps36^#25^*^86^), *Syntaxin 7* (*Syx7^#31^*^19^), *Rabaptin-5-associated exchange factor for Rab5* (*Rabex-5^#36^*^72^), *Tumor susceptibility gene 101* (*TSG101^#36^*^22^; later replaced by *TSG101^P^*^26^ ^43^ due to lethality), *furry* (*fry^#37^*^92^), *fmt^#33^*^55^ ^35^, *emei^#566h^* ^36^, and *msn^#32^*^08^ ^38^ (Fig. S1A). Disruption of these genes through induction of homozygous mutant clones in *Ras^V12^*cells led to enhanced tumor overgrowth and delayed pupariation of third instar larvae (Figs. S1B-E). These phenotypic effects mirrored those observed in *Ras^V12^* cells with disruptions in canonical cell polarity genes, such as *scrib^1^* ^27^ and *l(2)gl^4^* ^44^ (Figs. S1B-E). Interestingly, by dissecting larvae bearing *Ras*-driven tumors, we also observed tumor metastasis in the intestinal tissues (Figs. S1F), indicating that these *Ras*-driven tumors may possess the potential for further malignant progression. Thereby, summarizing these tumor suppressors, we obtained a cohort of ten *Drosophila Ras*-driven tumor (DRT) models.

### The Atlas of *Ras*-driven Tumors in *Drosophila* (ART-D)

Modeling tumorigenesis is crucial for deepening our understanding of cancer. To this end, we systematically characterized ten DRTs by sequentially tracking the tumorigenesis process through two complementary lenses: phenotypic malignancies and transcriptome dynamics (Fig. 1A). Phenotypic malignancies were evaluated using four key metrics: larval tumor burden, tumor-mediated cachexia, tumor overgrowth, and neighboring tissue invasion (Fig. 1A). We designated three time points (5.5 days after egg laying [AEL], 9.5 AEL, and 13.5 AEL) to collect phenotypic data and transcriptomic data (Fig. 1A), forming the Atlas of *Ras*-driven Tumors in *Drosophila* (ART-D) (Fig. 1B). Through ART-D, we found all tumors exhibited increasingly aggressive phenotypes over time and staged characteristics: 1) Initial stage: (0–5.5 days AEL): Marked by tumor cell overgrowth. 2) Progression stage, (5.5–9.5 days AEL): Characterized by uncontrolled proliferation, onset of invasion, and early cachexia. 3) Advanced stage (9.5–13.5 days AEL): Distinguished by severe cachexia and aggressive tissue invasion.

**Figure 1.**
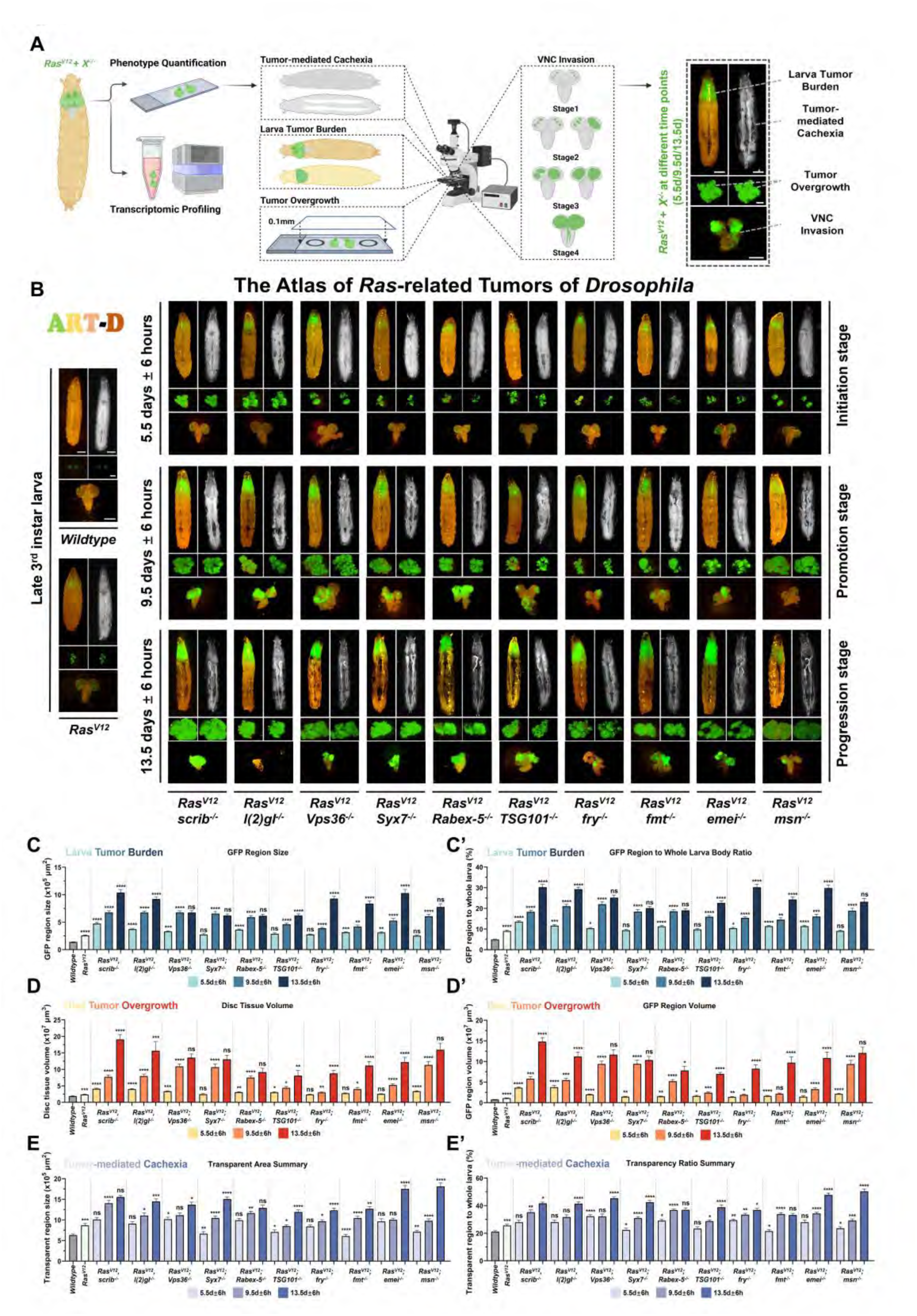
**The Atlas of *Ras*-driven Tumors of *Drosophila* (ART-D).**(A) The schematic diagram outlining the protocol for harvesting malignant tumor samples at designated time points (5.5d ± 6h, 9.5d ± 6h, and 13.5d ± 6h AEL) to assess phenotypic data across four categories: larva tumor burden, tumor-mediated cachexia, tumor overgrowth, and ventral nerve cord (VNC) invasion. Scale bars, 500 µm for larva tumor burden and tumor-mediated cachexia, 400 µm for tumor overgrowth, and 400 µm for VNC invasion. (B) The Atlas of *Ras*-driven Tumors of *Drosophila* (ART-D). The summary of four representative phenotypic characteristics of normal tissue, *Ras* benign tumors, and ten malignant tumors at designated time points. Three stages with distinct phenotypic features were identified in ten *Drosophila Ras*-driven malignant tumors: initial stage, promotion stage, and progression stage. Scale bars: 500 µm for larva tumor burden and tumor-mediated cachexia, 400 µm for tumor overgrowth, and 400 µm for VNC invasion. (C-E’) The quantifications of larval tumor burden, tumor-mediated cachexia, and tumor overgrowth in ART-D. The larval tumor burden was quantified by the GFP region size (C) and the ratio of GFP region to whole larva (C’); the tumor overgrowth was quantified by the overall disc tissue volume (D) and the GFP region volume (D’); and the tumor-mediated cachexia was quantified by the transparent region size (E) and the ratio of transparent region to whole larva (E’), respectively. Statistical analysis for each group was performed using the Mann-Whitney U-test and presented as mean ± SEM. ns > 0.05, \**P* < 0.05, \*\**P* < 0.01, \*\*\**P* < 0.001, \*\*\*\**P* < 0.0001, indicating the differences in larva tumor burden, tumor-mediated cachexia, or tumor overgrowth at the current stage compared to the previous stage, *Ras* group was compared with *wildtype* group, the initial stage tumors were compared with *Ras* group.

Larval tumor burden, quantified via GFP signal intensity and tumor-to-body ratio, reflected overall progression across genetic backgrounds. *Ras^V12^* tumors with disruptions in *scrib*, *l(2)gl*, *TSG101*, *fry*, *fmt*, or *emei* showed steadily increasing burden from initial to progression stages. In contrast, tumors lacking *Vps36*, *Syx7*, *Rabex-5*, or *msn* peaked in burden during the progression stage. It was noteworthy that the malignant tumors induced by the loss of *fry*, *fmt*, or *emei* exhibit a tumor burden at advanced stages comparable to that resulting from the loss of cell polarity genes (*scirb* and *l(2)gl*) (Figs. 1C and 1C’). Tumor overgrowth, measured by volumetric analysis of whole tissues and the GFP-labeled regions, mirrored these trends: *scrib*, *l(2)gl*, *TSG101*, *fry*, *fmt*, and *emei* mutants exhibited dramatic expansion from progression to advanced stages, while *Vps36*, *Syx7*, *Rabex-5*, and *msn* mutants showed maximal proliferation rates earlier, during the progression stage (Figs. 1D and 1D’). These results suggested that although ten DRTs exhibit malignant progression phenotypes, their growth rates differ.

Cachexia worsened progressively due to muscle and fat body degradation ^39, 41^, served as a hallmark feature of tumors stepped into promotion stage (Fig. 1B). All DRTs exhibited significant increase in either transparent area or transparency ratio at promotion stage, except for *Vps36* loss (Figs. 1E and 1E’). Moreover, *Ras^V12^* tumors lacking *l(2)gl*, *Vps36*, *Syx7*, *TSG101*, *fry*, *emei*, or *msn* all exhibited more drastic increase in both tumor-mediated cachexia parameters at progression stage (Figs. 1E and 1E’). Quantitative results suggested that tumor-induced cachexia begins to manifest during promotion stage of DRTs and progressively worsens as the tumor advances. Besides, the ventral nerve cord (VNC) was taken as neighboring tissue for invasion event quantification. Tumors with polarity defects (*scrib* or *l(2)gl* loss) displayed early invasion (25% stage 4 cells at initial stage), increasing over time. Endocytosis-deficient tumors (*Vps36*, *Syx7*, *Rabex-5*, *TSG101*) showed severe invasion (>50% stage 4 cells) by the progression stage. *fry*-, *fmt*-, *emei*-, and *msn*-deficient tumors exhibited delayed invasion, peaking only in advanced stages, with *msn* mutants showing weakest invasively (Figs. S1G). Collectively, the ART-D comprehensively delineated the phenotypic characteristics of *Ras^V12^* malignant tumors in stages, providing phenotypic references for elucidating the underlying molecular mechanisms.

### Three Subtypes of *Drosophila Ras*-driven Tumors

Apart from four malignancy traits, we also found DRTs exhibited general characteristics of malignant tumor models, strong autonomously overgrowth and genome instability at the initial stage of tumorigenesis, as revealed by immunofluorescent staining of PH3 and γH2AV (Fig. 2A and 2B). However, the quantification of malignancies revealed intra-tumor heterogeneity among DRTs, indicating the progression trajectories diverged from different tumor models. To explore the preferences of different tumor models on phenotypic malignancies, we used robust linear regression to six digitalized phenotypic malignancies: GFP region size and GFP region to whole larva body ratio (Fig. S2A and S2A’), disc tissue volume and GFP region volume (Fig. S2B and S2B’), transparent area summary and transparency ratio summary (Fig. S2C and S2C’).

**Figure 2.**
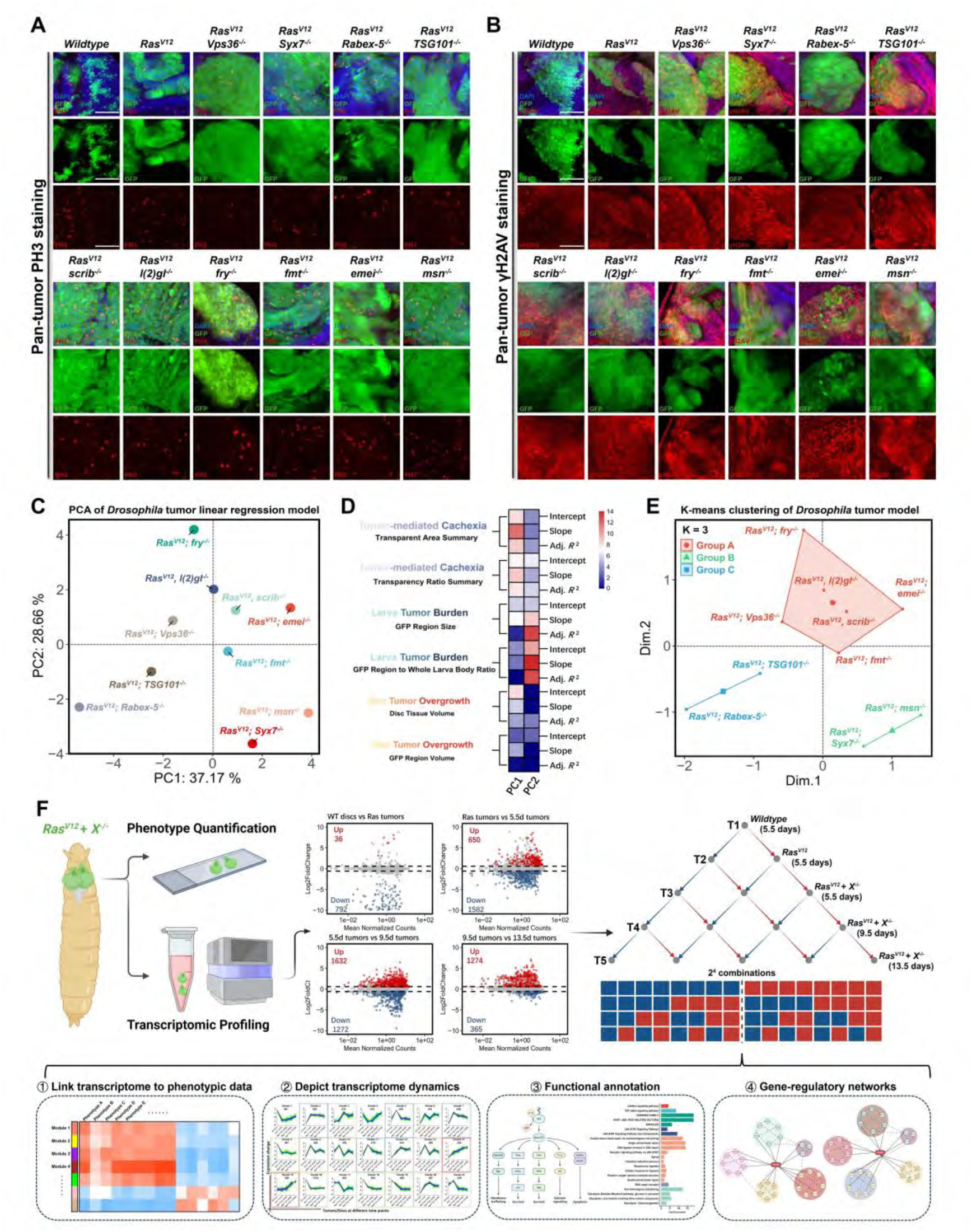
**The classifications of *Drosophila Ras*-driven tumors.** (A) Confocal images of eye-antennal discs or tumors harvested at 5.5-6.5 days AEL. Discs or tumors with indicated genotypes were stained with antibody against PH3. Scale bars: 50 µm. (B) Confocal images of eye-antennal discs or tumors harvested at 5.5-6.5 days AEL. Discs or tumors with indicated genotypes were stained with antibody against γH2AV. Scale bars: 50 µm. (C) The two-dimensional scatter plot visualizing the PCA projection of *Drosophila Ras*-driven tumors. The PCA was performed based on the robust linear regression model parameters of each *Ras*-driven tumor, including the intercept, slope, and adjusted R-squared values. (D) The contributions of PC1 and PC2 to the intercept, slope, and adjusted R-squared values of robust linear regression models representing tumor malignancies. (E) The K-means clustering of *Drosophila Ras*-driven tumors based on PCA results. A total of three groups were classified, group A comprises *scrib*-, *l(2)gl*-, *fry*-, *Vps36*-, and *emei*-deficient tumor models; group B comprises *Syx7*- and *msn*-deficient tumor models; group C comprises *Rabex5*- and *TSG101*-deficient tumor models. (F) The schematic diagram outlining the workflow for identifying tumor transcriptomic dynamics. The process began with the acquisition of bulk RNA-seq data from various tumors at different time points (Step 1), followed by the identification of DEGs (Step 2), the analysis of gene dynamic changing patterns (Step 3), and the extension of analyses (①∼④) link transcriptomic changes with phenotypic malignancies (Step 4).

The intercept was largely affected by the malignancies at the initial stage, while the slope indicated the progression rate from initial stage to progression stage. Comparing the slopes of each regression equation with average regression equation, we found the *scrib*-, *l(2)gl*-, *fry*-, *emei*-, and *msn*-deficient tumors exhibited more aggressive progression on larva tumor burden than the average levels of all tumors, 0.59 for GFP region size and 1.81 for GFP region ratio (Fig. S3A and S3A’). Disruptions of *scrib*, *Syx7*, *emei*, or *msn* exhibited more aggressive progression on tumor overgrowth, 1.11 for disc tissue volume and 1.01 for GFP region volume (Fig. S3B and S3B’). *Ras^V12^* tumors with *Syx7*, *emei*, or *msn* loss were also found higher progression level in tumor-mediated cachexia manifestations compared to the average progression level of all tumors, 0.732 for transparent area and 1.85 for transparency ratio (Fig. S3C and S3C’). The varying degrees of progression in different tumors prompted us to consider whether tumors can be classified based on these characteristics.

Using the intercepts, slopes, and adjusted R-squared values from robust linear regression models of each DRT as input, we conducted principal component analysis (PCA) to project the tumor models onto a two-dimensional map (Fig. 2C). The contributions of principle components (PC1 and PC2) to malignancies were shown, with PC1 capturing more features associated with tumor-mediated cachexia and PC2 reflecting more features related to larval tumor burden (Fig. 2D). Building on the PCA results, we applied K-means clustering for tumor subtyping. Consequently, Group A included *Ras^V12^* tumors with deficiencies in *scrib*, *l(2)gl*, *Vps36*, *fry*, *emei*, or *fmt*; Group B consisted of tumors lacking *Syx7* or *msn*; and Group C contained tumors deficient in *TSG101* or *Rabex-5* (Fig. 2E).

### Transcriptome Dynamics of *Drosophila Ras*-driven Tumor Subtypes

To elucidate the molecular mechanisms driving malignant transformation across different tumor subtypes, we established five pseudotime points to simulate longitudinal progression from initial to advanced stages: T1 (wild type at 5.5 days AEL), T2 (*Ras^V12^* at 5.5 days AEL), T3 (*Ras^V12^,X^-/-^* at 5.5 days AEL), T4 (*Ras^V12^,X^-/-^*at 9.5 days AEL), and T5 (*Ras^V12^,X^-/-^* at 13.5 days AEL). Theoretically, five pseudotime points encompassed 16 (2^4^) possible combinations of gene expression dynamic change patterns (Fig. 2F). Following batch effect correction (Fig. S4A and Supplementary Table_S1), we notified the sustained upregulation (clusters 1, 4, 14, 15) or downregulation (clusters 9, 19, 21) trends were the dominant patterns of all DRTs (Fig. S4B). However, not all observed dynamic changes were closely associated with the tumor malignancy progression. Thereby, we conducted integrative analyses of phenotypic and transcriptomic data across different tumor subtypes to reveal dynamic transcriptomic features associated with malignant tumor progression (Fig. 2F).

In Group A DRTs, weighted gene co-expression network analysis (WGCNA) ^45^ identified seven gene modules. Among these, the MEbrown and MEred gene modules showed significant positive correlations with multiple malignancies, whereas MEblue and MEturquoise gene modules exhibited significant negative correlations (Fig. 3A). By extracting genes from these modules and intersecting them with differentially expressed genes (DEGs, T2 vs T5, Supplementary Table_S2), we identified 1,454 upregulated and 1,583 downregulated DEGs significantly associated with tumor malignancy progression (Fig. 3B and 3C). Subsequently, Mfuzz ^46^ clustering was applied to delineate the transcriptomic dynamics of these genes, revealing continuously upregulated gene clusters (Clusters 4, 5, and 8) and continuously downregulated gene clusters (Clusters 1, 2, 3, 6, and 7) from stage T1 to T5, which were selected as candidate gene sets (Fig. 3D). Functional annotation using PANGEA indicated that, in Group A tumors, the biological processes significantly associated with phenotypic malignancies primarily included glycolysis, glyconeogenesis, pentose phosphate pathway, amino acid synthesis, and fatty acid metabolism. The signaling pathways significantly linked to phenotypic malignancies mainly involved the Notch, EGFR/FGFR/PvR, and TNFɑ signaling pathways (Fig. 3E).

**Figure 3.**
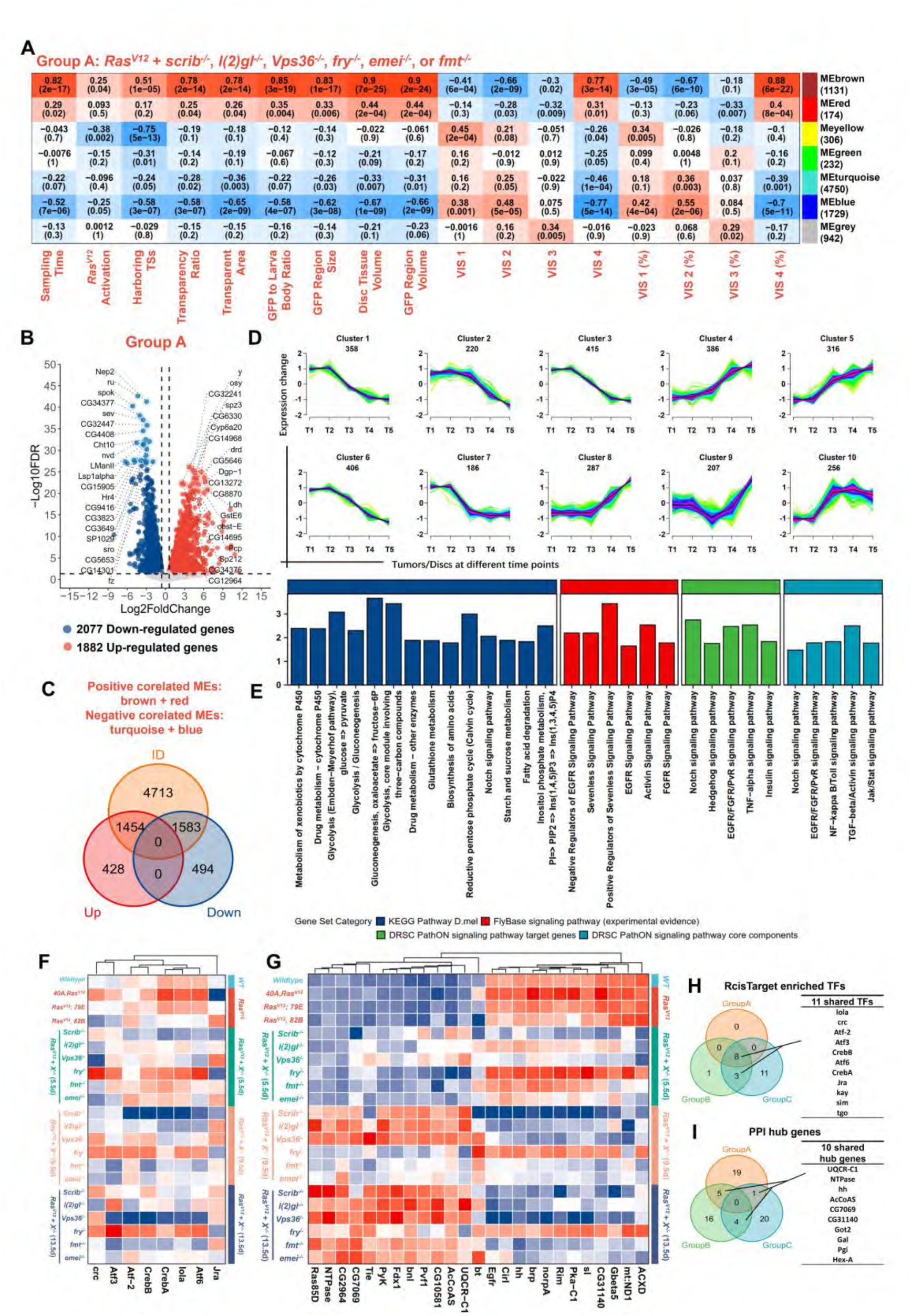
**Dynamic transcriptomic profiles of Group A *Drosophila Ras*-driven tumors.** (A) The WGCNA heatmap depicting the correlations between identified gene modules and phenotypic malignancies of Group A *Drosophila Ras*-driven tumors. The color intensity of each module represented the correlation coefficient, which was also displayed in the top line of the respective cell. The corresponding *P*-value was shown in parentheses, with *P* values below 0.05 indicating statistically significant correlations. (B) The volcano plot of DEGs (T2 vs T5) in Group A of *Drosophila Ras*-driven tumors. The up-regulated and down-regulated genes were 1,882 and 2,077, respectively. (C) The Venn diagram depicting the overlap of genes in modules significantly associated with tumor malignancies (A) and DEGs (B). 1,454 genes were up-regulated and significantly associated with tumor malignancies; 1,583 genes were down-regulated and significantly associated with tumor malignancies. (D) The line charts depicting dynamic transcriptomic changes of DEGs significantly associated with tumor malignancies in Group A *Drosophila Ras*-driven tumors (C), revealed by Mfuzz. (E) The bar plot depicting the functional annotations of candidate genes obtained from certain clusters (D), revealed by PANGEA. (F) The heatmap depicting the expression profiles of enriched TFs from T1 to T5. The TF enrichment of candidate genes derived from specific clusters (D) was performed using RcisTarget. (G) The heatmap depicting the expression profiles of identified hub genes from T1 to T5. The hub gene identification among candidate genes from specific clusters (D) was performed via STRING and BottleNeck. (H) The Venn diagram depicting the overlap of shared TFs in different groups of *Drosophila Ras*-driven tumors. The shared TFs were listed on the table. (I) The Venn diagram depicting the overlap of shared hub genes in different groups of *Drosophila Ras*-driven tumors. The shared hub genes were listed on the table.

By dissecting the transcriptome dynamics of Group B and Group C DRTs, we found the functional annotation of continuously upregulated/downregulated gene clusters significantly associated with tumor malignant progression in Group B primarily involve amino acid synthesis, glycolysis and gluconeogenesis, fatty acid oxidation, synthesis of unsaturated fatty acids, and triglyceride biosynthesis. The signaling pathways significantly associated with tumor malignant progression mainly include EGFR/FGFR/PvR, JAK/STAT, and Hippo pathways (Fig. S5A-E). In Group C DRTs, the biological processes significantly associated with tumor malignant progression, apart from glycolysis and gluconeogenesis, also include inositol phosphate metabolism and O-glycan biosynthesis. The signaling pathways significantly associated with tumor malignant progression mainly include TGFβ/Activin, EGFR/FGFR/PvR, and Notch pathways (Fig. S6A-E).

By horizontally comparing the different tumor subtypes, we found the common features across tumor subtypes are enriched in biological processes such as glycolysis, gluconeogenesis, and amino acid metabolism, as well as the EGFR/FGFR/PvR signaling pathway, indicating the common characteristics for maintaining tumor malignant progression during T1 to T5 of DRTs. Meanwhile, we also found a large number of biological processes related to fatty acid metabolism enriched in Group B, while biological processes related to inositol phosphate metabolism and O-glycan biosynthesis were enriched in Group C, revealing the specific metabolic features of tumor Groups B and C in maintaining tumor malignant progression from T1 to T5. Notably, in terms of signal transduction, both tumor

Groups A and C were enriched in Notch signaling pathway and NF-κB/Toll signaling pathway, while tumor Groups A and B were enriched in the JAK/STAT signaling pathway. The Hippo signaling pathway was unique to Group B, and the TGFβ/Activin signaling pathway showed higher enrichment in Group C.

Through transcription factor enrichment analysis and hub gene identification of DEGs significantly associated with tumor malignant progression in different tumor subtypes, we found that Notch signaling pathway-related transcription factor *longitudinals lacking* (*lola*), as well as the MAPK signaling pathway-related transcription factors *Jun-related antigen* (*Jra*) and *kayak* (*kay*), were co-enriched in different tumor groups (Figs. 3F and 3H, Fig. S5F, and Fig. S6F). Based on the protein-protein interaction networks established using STRING database^47^, *Glutamate oxaloacetate transaminase 2* (*Got2*), *NTPase*, *Phosphoglucose isomerase* (*Pgi*), and *Hexokinase A* (*Hex-A*) were identified as hub genes in at least two tumor subtypes (Figs. 3G and 3I, Fig. S5G, and Fig. S6G). These results suggest that the biological processes or signaling pathways regulated by these transcription factors and hub genes may play important roles in maintaining tumor malignant progression.

### Activation of NF-κB/Toll and MAPK signaling in T3 stage across all tumor subtypes

To further investigate signaling pathways that were crucial for tumor initiation, we compared different subtypes of DRTs (T3) with *Ras* benign tumors (T2). The intersection showed 247 co-upregulated genes and 578 co-downregulated genes in DRTs at the T3 stage (Fig. S7A). Functional annotation revealed that these genes were primarily involved in regulating the JAK/STAT, NF-κB/Toll, and MAPK signaling pathways (Fig. S7B), consistent with the common features observed in the dynamic transcriptomic profiles of different tumor subtypes from T1 to T5. Additionally, we found that the expression levels of JAK/STAT ligands *Unpaireds* (*Upds*: *Upd1*, *Upd2*, and *Upd3*), the receptor *Toll* (*Tl*) and the transcription factor *dorsal* (*dl*) from the NF-κB/Toll signaling pathway, as well as the JNK signaling pathway target genes *MMP1* and *puckered* (*puc*), were higher at the T3 than T2, with expression levels increasing progressively during tumor development (Fig. S7C). Together, these findings suggest that the JAK/STAT, NF-κ B/Toll, and MAPK signaling pathways may be simultaneously activated in various tumor subtypes at the T3 stage.

Bulk-level dynamic gene expression analyses may overlook tumor-specific characteristics. To address this, we used gene set variation analysis (GSVA) ^48^ and single-sample gene set enrichment analysis (ssGSEA) ^49, 50^ to score 15 major signaling pathways in *Drosophila* (Fig. S7D and S7E). To minimize interference from antagonistic effects, we focused on positive regulators of signaling pathways to evaluate pathway activation across ten *Drosophila* tumor models from T1 to T5 (Fig. S7D). Results showed consistent activation of JAK/STAT signaling from T3 to T5 in eight genotypic tumors, with activation in all genotypic tumors at the T5. Similarly, TNFɑ signaling was activated at the T3 stage in nearly all tumors. Other MAPK-related signaling pathways (EGFR, FGFR, PvR, Sevenless, and Torso) were primarily activated at T3, although FGFR and Torso signaling were inactivated in endocytosis-impaired tumors (*Vps36⁻/⁻*, *Syx7⁻/⁻*, *Rabex-5⁻/⁻*, and *TSG101⁻/⁻*). The widespread activation of JAK/STAT and MAPK signaling since or at T3 in DRTs further elucidates the positive correlations between these two pathways and tumor progression at the bulk-level of transcriptome dynamics.

Notably, *l(2)gl*-, *TSG101*-, *fry*-, or *fmt*-deficient tumors exhibited co-activation of the NF-κB/Toll and NF-κB/Imd pathways at T3. Insulin signaling was activated at T3 in seven tumor types and remained continuously active from T1 to T5 in *Ras^V12^* tumors with disrupted *l(2)gl*, *TSG101*, *emei*, *fmt*, or *msn*. Given that JNK signaling activation can lead to increased transcription the ligands of JAK/STAT signaling *Upds*, we selected the JNK and NF-κB signaling pathways for experimental validation. Immunofluorescence staining revealed increased phosphorylation levels of the MAPK pathway effectors *rolled* (*rl/ERK*) and *basket* (*bsk/JNK*), as well as elevated protein levels of their downstream targets *MMP1* and *wingless* (*wg*), in all DRTs (Fig. S8A-D), consistent with our hypothesis of extensive MAPK pathway activation. Furthermore, immunofluorescence staining of the key downstream transcription factors *dl* and *Relish* (*Rel*) of NF-κB/Toll and NF-κB/Imd pathways showed consistently elevated dl protein levels across all DRTs, whereas Rel expression increased only in tumors with specific genotypes, such as *Ras^V12^* tumors with *fmt* deficiency (Fig. S8E and S8F). Collectively, these findings suggest that activation of the MAPK and NF-κB/Toll signaling pathways may represent common mechanisms underlying the malignant progression of DRTs.

### Notch Activation and Hippo Inactivation Facilitate Excessive Tumor Overgrowth

To further assess the oncogenic potential of major signaling pathway dynamics, we correlated pathway activation status with tumor progression parameters. Notably, Notch signaling activation exhibited a positive correlation with tumor progression, whereas Hippo signaling activation inversely correlated with severity across DRTs (Figs. 4A, 4B, S9A, and S9B). To validate the broad dysregulation of these pathways in DRTs, we performed immunofluorescent staining of key Notch components, including the ligand *Delta* (*Dl*) and Notch intracellular domain (*NICD*), and its target *dMyc*, and the transcriptional co-factor of the Hippo pathway, *Yorkie* (*Yki*), and its downstream target gene *Death-associated inhibitor of apoptosis 1* (*Diap1*). Compared to wild type discs and *Ras* benign tumors, robust upregulation of Delta, NICD, dMyc, and Diap1, as well as increased nuclear translocation of Yki (Figs. 4A, 4D, and S9C-E), was observed across tumor samples, confirming Notch hyperactivation or Hippo inactivation.

**Figure 4.**
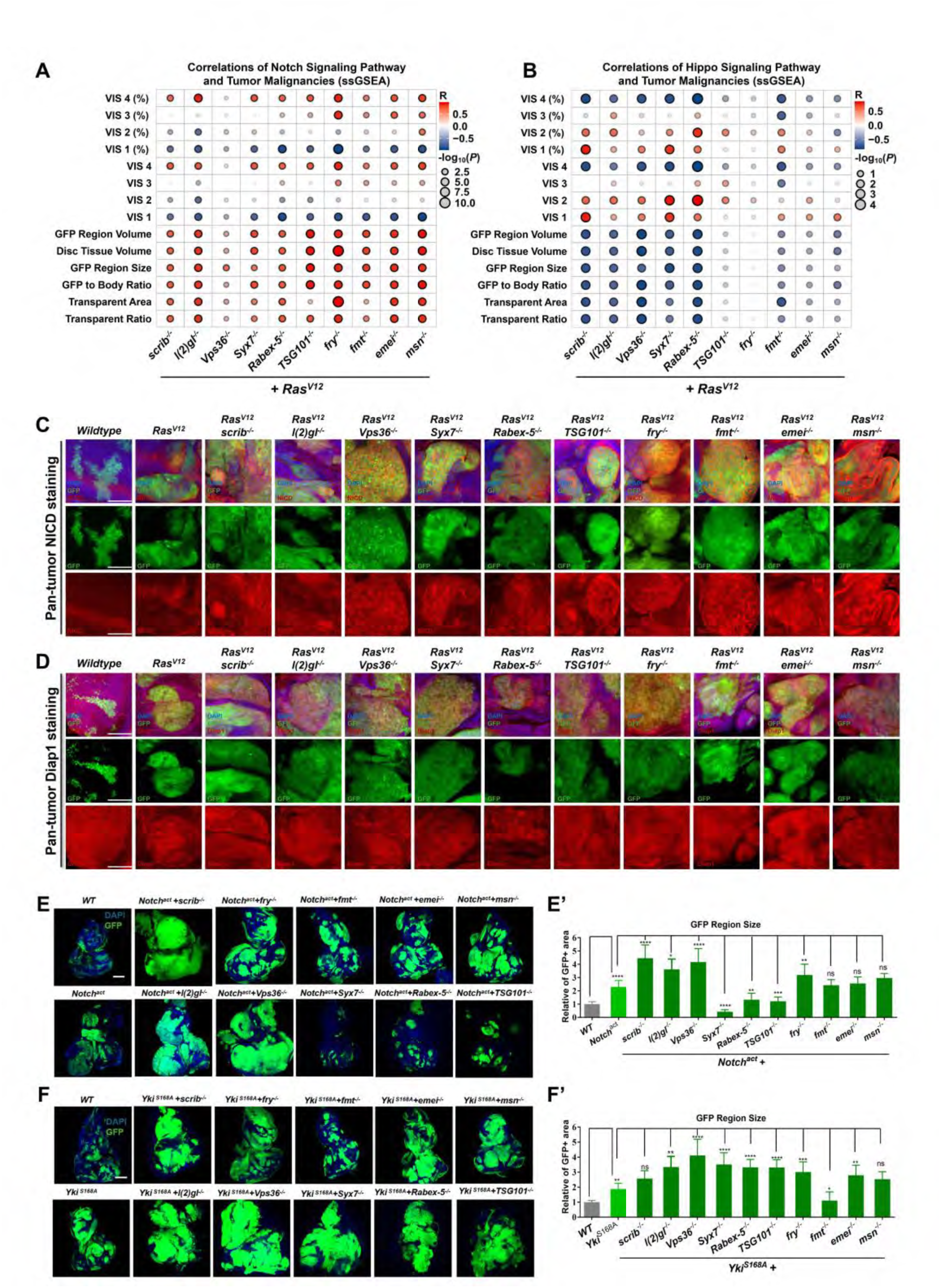
**The tumor promoting role of Notch activation and Hippo inactivation within of *Drosophila Ras*-driven tumors.** (A) The dot plot depicting the correlations between Notch signaling pathway scores and tumor malignancies. The signaling scores across ten *Drosophila Ras*-driven tumors were evaluated by ssGSEA. The correlations were evaluated by Pearson correlation test. The color of the dots indicated the Pearson correlation coefficients (R), while the size of the dots reflected the negative log10-transformed *P* values. (B) The dot plot depicting the correlations between Hippo signaling pathway scores and tumor malignancies. The signaling scores across ten *Drosophila Ras*-driven tumors were evaluated by ssGSEA. The correlations were evaluated by Pearson correlation test. The color of the dots indicated the Pearson correlation coefficients (R), while the size of the dots reflected the negative log10-transformed *P* values. (C-D) Confocal images of eye-antennal discs or tumors harvested at 5.5-6.5 days AEL. Discs or tumors with indicated genotypes were stained with antibodies against Notch and Hippo signaling pathway members, NICD (C) and Diap1 (D), respectively. Scale bars: 50 µm. (E-E’) The confocal images (E) and quantifications of GFP region size (E’) within discs or tumors overexpressing the activated form of *Notch* (*Notch^act^*) along with corresponding tumor suppressors. Statistical analysis for each group was performed using the Mann-Whitney U-test as indicated and presented as mean ± SD. ns > 0.05, \**P* < 0.05, \*\**P* < 0.01, \*\*\**P* < 0.001, \*\*\*\**P* < 0.0001. (F-F’) The confocal images (F) and quantifications of GFP region size (F’) within discs or tumors overexpressing the activated form of *Yki* (*Yki^S^*^168^*^A^*) along with corresponding tumor suppressors. Statistical analysis for each group was performed using the Mann-Whitney U-test as indicated and presented as mean ± SD. ns > 0.05, \**P* < 0.05, \*\**P* < 0.01, \*\*\**P* < 0.001, \*\*\*\**P* < 0.0001.

Next, functional validation of Notch and Hippo signaling in tumorigenesis was conducted by expressing constitutively active *Notch* (*Notch^act^*) or *Yki* (*Yki^S168A^*) in GFP-marked cells deficient for specific tumor suppressors. *Notch^act^* synergized with the loss of *scrib*, *l(2)gl*, *Vps36*, or *fry* to drive aggressive tumor overgrowth, characterized by giant larvae phenotypes in *scrib*- and *Vps36*-deficient tumors (Figs. 4E, 4E’, S9F, and S9H). Conversely, *Notch^act^* failed to promote clone expansion in cells lacking *Syx7*, *Rabex-5*, or *TSG101* (Figs. 4E, 4E’, S9C, and S9H). The discrepancy of *Notch^act^* synergizing endocytosis-defects may be due to the interference of different endocytosis processes. Previous studies showed defects in *Vps36*, *Syx7*, or *TSG101* could drive tumorigenesis and exhibit elevated Notch due to internalization, but the target gene expression only elevated in *TSG101*-deficient cells ^51–53^. Similarly, *Yki^S168A^* cooperated with defects in *l(2)gl*, *Vps36*, *Syx7*, *Rabex-5*, *TSG101*, *fry*, or *emei* to promote tumor overgrowth, inducing giant larvae phenotypes in polarity-deficient cells (Figs. 4F, 4F’, S9G, and S9I). These results imply a context-dependent synergy between tumor suppressor loss and hyperactivated *Notch* or *Yki*. Intriguingly, we noticed that *l(2)gl*-, *Vps36*-, and *fry*-deficient cells exhibited unique potency, cooperating with multi oncogenic mutations (*Ras^V12^*, *Notch^act^*, or *Yki^S168A^*) to fuel malignancy (Figs. S1B-E and 4E-F’), suggesting that these potent tumor suppressors may converge on shared mechanisms enabling tumor plasticity.

### Multi-omics Analysis Links JNK/Notch/Toll Activation and Hippo Inactivation to Tumor Progression

Among these three tumor suppressors, *fry* encodes a 3,479-amino acid protein that functions in a complex with the NDR (Nuclear Dbf2-driven) kinase tricornered (trc) and mob as tumor suppressor (mats) ^54–56^. Fry-NDR kinase signaling is implicated in regulating polarized cell growth and epidermal morphogenesis, with *fry* and *trc* mutants exhibiting increased cell volume ^54, 57^. Notably, depletion of *FRY* could promote the nuclear localization of YAP/TAZ with decreased NDR1/2 kinase activities and YAP phosphorylation levels independent of LATS1/2 activity ^58^. Moreover, downregulation of the *trc* ortholog, *NDR1*, promotes epithelial-mesenchymal transition (EMT) in prostate cancer, while its overexpression suppresses metastatic phenotypes ^59^. These findings suggest a conserved tumor-suppressive role for *fry* in oncogenic contexts, though the underlying mechanisms remain unclear.

To address this gap, we performed an integrative multi-omics analysis of bulk RNA-seq and Assay for Transposase-Accessible Chromatin with high-throughput sequencing (ATAC-seq) on *Ras^V12^; fry*^−/−^ tumors, a novel model previously uncharacterized in cancer studies, across distinct progression stages (Fig. 5A). Initial DEG analysis revealed no sustained up- or downregulated candidates upon cross-stage comparison (Fig. S10A), we subsequently employed WGCNA to identify dynamic transcriptional modules that positively correlated with tumor progression metrics (e.g., tumor burden, cachexia severity, VNC invasion; Figs. S10B–F). Module gene re-clustering and pathway annotation highlighted enrichment of MAPK, NF-κB/Toll, Notch, and Hippo signaling pathways (Figs. S10F, and S10G), suggesting their involvement in driving *Ras^V12^; fry^-/-^* tumor malignancies. Temporal expression profiling revealed stage-specific activation: *Tl*, *mastermind* (*mam*), *Diap1*, *frizzled* (*fz*), and *wg* peaked at T3, while Insulin signaling genes (*GlyP*, *Lpin*, *Ilp2*, *Ilp3*, *Thor*) were mainly upregulated at T4-T5, coinciding with aggressive cachexia and invasion (Fig. S10F). Additionally, ATAC-seq confirmed elevated chromatin accessibility at promoter regions of key effector/target genes in JNK, Hippo, NF-κB/Toll, and Notch pathways across tumor stages (Fig. 5B). Motif enrichment analysis of differentially accessible peaks identified top-ranked transcription factor binding sites for Initiator, DREF, and AP-1 (Fig. 5B). Collectively, our integrative analysis implicates JNK, NF-κ B/Toll, Hippo, and Notch signaling as cooperative drivers of tumorigenesis in *Ras*-activated, *fry*-deficient contexts.

**Figure 5.**
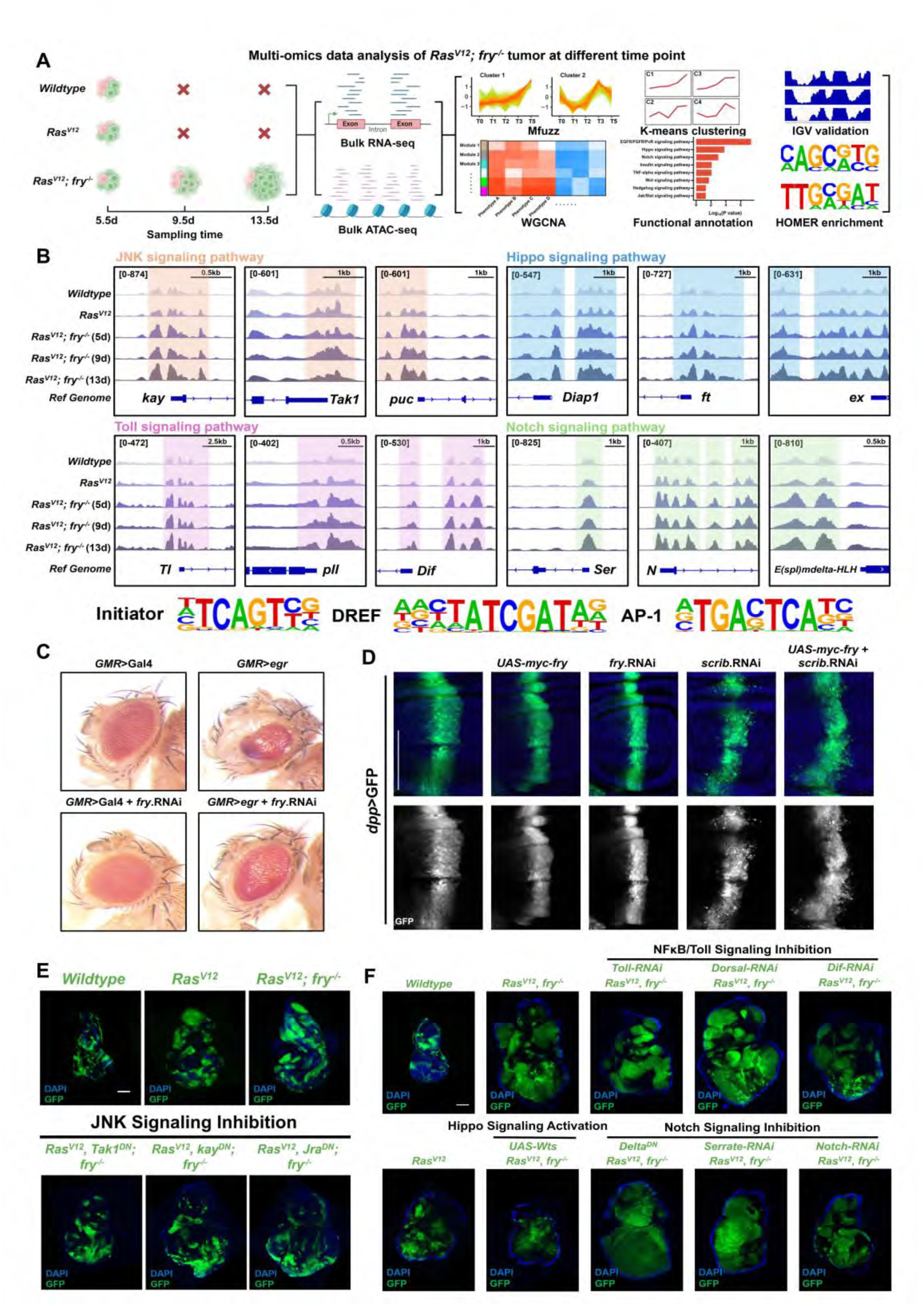
**Multi-omics data analysis of *Ras^V12^; fry^-/-^* tumors across different stages.** (A) The flowchart depicting the multi-omics data analysis of *Ras^V12^*; *fry^-/-^* tumors at various stages. Samples with indicated genotypes were harvested at designated time points (T1 to T5) for bulk RNA-seq and bulk ATAC-seq. Subsequent data analysis was conducted as outlined to uncover the mechanisms underlying tumorigenesis. (B) The IGV plot (top) illustrates the accessibility of the key effectors or target genes from specific signaling pathways, JNK signaling pathway (light orange), Notch signaling pathway (light green), Hippo signaling pathway (light blue), and Toll signaling pathway (light magenta). Data were normalized and presented as RPKM, with the maximum signal intensity and the scale bar for each region were denoted at the top. The identified top-ranked transcription factor binding motifs and their corresponding transcription factors (bottom). (C) The light micrographs of adult eyes with the indicated genotypes. (D) The fluorescence micrographs of wing pouch regions, wing discs were placed as anterior to the left and dorsal up. Scale bars: 100 µm. (E) The confocal images of JNK signaling inhibition in *Ras^V12^; fry^-/-^* tumors. Samples were observed at 5.5d AEL. Scale bar: 100 µm. (F) The confocal images of NF-κB/Toll signaling inhibition, Hippo signaling inactivation, and Notch signaling inhibition in *Ras^V12^; fry^-/-^* tumors as indicated. Samples were observed at 5.5d AEL. Scale bar: 100 µm.

To investigate the physiological and pathological roles of *fry* in regulating the JNK, NF-κ B/Toll, Hippo, and Notch signaling pathways, we knocked down *fry* expression in the posterior region of the *Drosophila* wing disc and examined the expression levels of target genes or key kinases associated with these pathways (Figs. S11A and S11B). No significant differences were observed compared to the *wildtype* controls. Similarly, *fry* knockdown in the *Drosophila* adult eye and thorax did not affect eye size or the JNK-dependent thorax closure process (Figs. 5C and S11C). However, under pathological conditions, inhibition of *fry* rescued the *eiger* (*egr*)-induced small eye phenotype (Fig. 5C), and overexpression of *fry* partially suppressed *scrib* knockdown-induced JNK-dependent cell migration (Fig. 5D). These findings suggest that while *fry* does not exert a significant regulatory influence on these pathways under physiological conditions, it can modulate JNK-dependent processes in pathological contexts.

To validate the functional necessity of these pathways in driving tumorigenesis, we inhibited JNK signaling in *Ras^V12^*; *fry^-/-^*tumors using dominant-negative forms of its key effectors (*TGF-β activated kinase 1* [*Tak1*], *kay*, and *Jra*). This intervention resulted in a marked reduction in imaginal disc size and rescue of the pupariation defects (Figs. 5E and S11D). Further investigation of the oncogenic roles of NF-κB/Toll, Notch, and Hippo pathways revealed that knocking-down of *Tl* and *Dorsal-driven immunity factor* (*Dif*) to suppress NF-κB/Toll signaling inhibited tumor growth, whereas *dl* knockdown had no effect (Fig. 5F). By inhibiting the activity of the Notch signaling pathway ligand *Delta*, and knocking down ligand *Ser* and receptor *Notch*, we found that inhibiting Delta did not significantly suppress tumor growth, while knocking-down *Ser* and *Notch* exhibited a trend of tumor growth inhibition (Fig. 5F). By overexpressing *warts* (*wts*) to activate Hippo signaling, we observed that the suppression of *Ras^V12^*; *fry^-/-^* tumors (Fig. 5F). Interestingly, we found that interfering with the activity of either signaling pathway could significantly rescue the pupariation delay defect in larvae bearing *Ras^V12^*; *fry^-/-^* tumors (Fig. S11E).

### Translational Analysis of *Drosophila* Tumor Suppressors in Human Cancers

To assess the clinical relevance of *Drosophila* tumor suppressors in human cancers with *RAS* activation, we analyzed pan-cancer genomic data obtained from cBioPortal ^60^ and gene expression data from UCSC-Xena^61^ (Fig. 6A). Among cancers with high *RAS* family mutation rates, *KRAS* mutations predominated in pancreatic adenocarcinoma (PAAD: 63.6%), colorectal adenocarcinoma (COADREAD: 37.4%), non-small cell lung cancer (NSCLC: 19.2%; LUAD: 31.8%; LUSC: 4.5%), uterine corpus endometrial carcinoma (UCEC: 20.0%), uterine carcinosarcoma (UCS: 17.5%), stomach adenocarcinoma (STAD: 16.4%), and esophageal carcinoma (ESCA: 8.8%). *NRAS* mutations were prevalent in skin cutaneous melanoma (SKCM: 28.4%) and thyroid carcinoma (THCA: 7.8%), while *HRAS* mutations were rare (<4%) (Fig. 6B). Based on mutation frequency, PAAD, COADREAD, LUAD, STAD, ESCA, LUSC, UCEC, and UCS were classified as *KRAS*-driven; SKCM and THCA as *NRAS*-driven (Fig. 6B).

**Figure 6.**
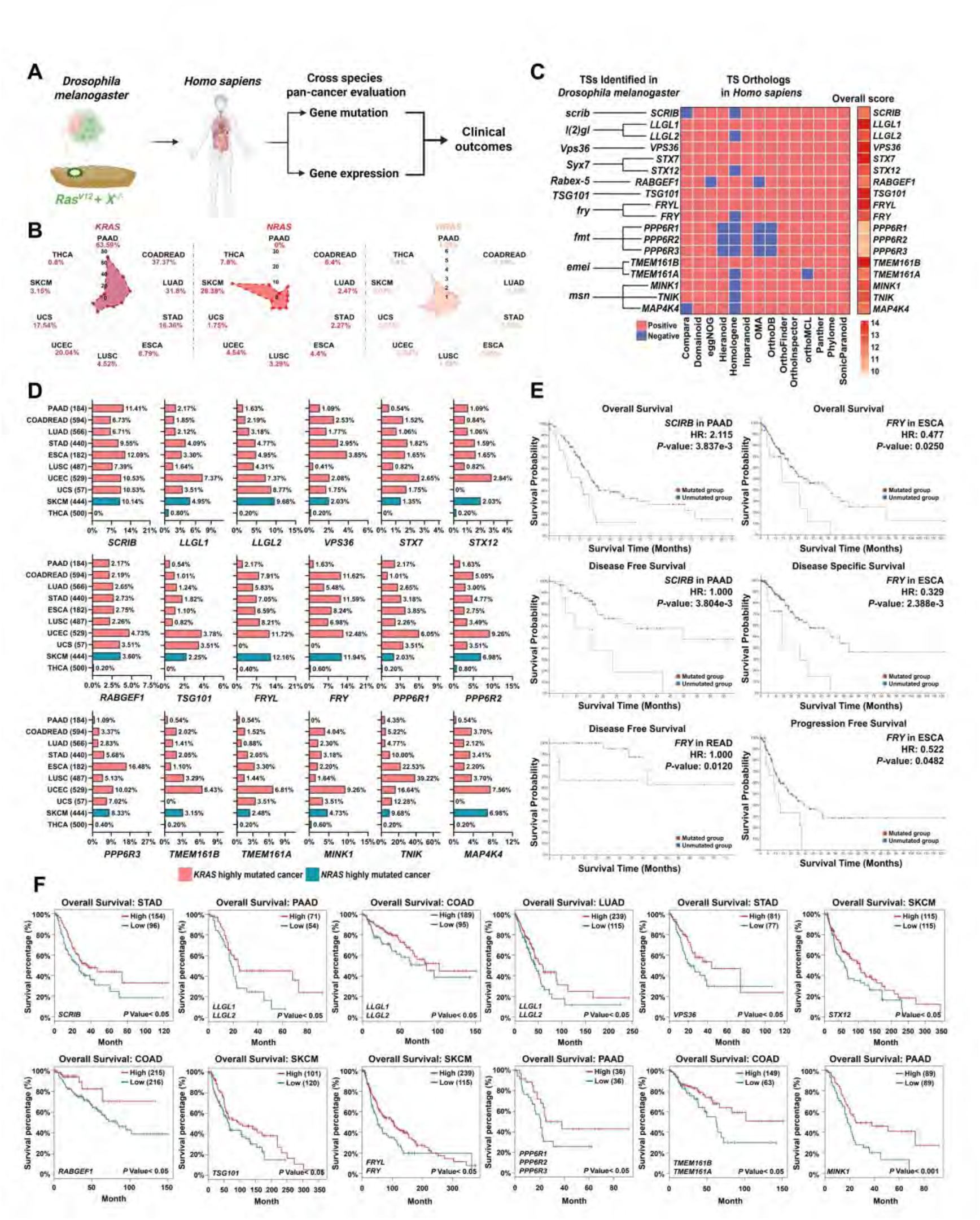
**The pan-cancer evaluation of human orthologs in *KRAS*/*NRAS*-driven cancers** (A) The scheme depicting the evaluation of the impact of *Drosophila* tumor suppressor human orthologs on human cancers from two perspectives, gene mutation and gene expression. (B) The mutation rate of *RAS* family genes (*KRAS, NRAS, HRAS*) in ten cancers (PAAD, COADREAD, LUAD, STAD, ESCA, LUSC, UCEC, UCS, SKCM, THCA). Data was obtained from TCGA PanCancer Atlas Studies of cBioProtal. (C) The eighteen *Homo sapiens* orthologs correspond to ten *Drosophila* tumor suppressors. The human orthologs were identified via 14 databases, the overall scores of each orthologous genes across different databases were presented in the right column. (D) The mutation rate of eighteen human orthologs across the ten cancer types that have high mutation rate of *KRAS* or *NRAS*. (E) Kaplan-Meier survival curves illustrating the impact of human ortholog mutation status on patient survival in specific cancers. (F) Kaplan-Meier survival curves illustrating the influence of human ortholog expression levels on overall survival in patients with *KRAS*/*NRAS*-driven cancers.

Using 14 public databases ^62–75^, we annotated these ten *Drosophila* tumor suppressors to 18 human orthologs: *scrib/SCRIB*; *l(2)gl*/*LLGL1*/*LLGL2*; *Vps36*/*VPS36*; *Syx7*/*STX7*/*STX12*; *Rabex-5*/*RABGEF1*; *TSG101*/*TSG101*; *fry*/*FRYL*/*FRY*; *fmt*/*PPP6R1*/*PPP6R2*/*PPP6R3*; *emei*/*TMEM161A*/*TMEM161B*; *msn*/*MINK1*/*TNIK*/*MAP4K4* (Fig. 6C). Mutation rates and expression levels of these orthologs were analyzed across *RAS*-driven cancers (Figs. 6D and S12A). *SCRIB* exhibited >10% mutation rates in PAAD, ESCA, UCEC, UCS, and SKCM, with downregulation in SKCM. *LLGL1*/*LLGL2* showed the highest mutation rates in UCEC (7.4%) and SKCM (9.7%), respectively, alongside reduced expression. Endocytosis-driven genes (*VPS36*, *STX7*, *STX12*, *RABGEF1*, *TSG101*) were frequently mutated in UCEC/SKCM and downregulated in these tumors, except *STX7* in SKCM. Notably, *FRYL* and *FRY* mutations exceeded 5% in most cancers (excluding PAAD, UCS, THCA), with *FRY* consistently downregulated across various cancers and co-decreased with *FRYL* in LUAD, LUSC, UCEC, and SKCM. These results suggest that the tumor suppressors identified in *Drosophila* model also exhibit high mutation rates and tumor suppressor potential in *KRAS*/*NRAS*-driven human cancers.

We then investigated the prognostic value of these human orthologs. In specific cancer types, single ortholog mutation is sufficient to confer a significant impact on the survival prognosis of patients (Fig. 6E). For instance, *SCRIB* mutations significantly threatened the overall survival (OS) and disease-free survival (DFS) of PAAD patients. Similarly, *FRY* mutations adversely affected the OS, disease-specific survival, and progression-free survival of ESCA patients. Besides, we observed significant positive correlations between the expression of human orthologs and the patient OS across various cancers (Fig. 6F). These findings suggest that the mutations and expression levels of these orthologs may serve as potential prognostic biomarkers in *KRAS*/*NRAS*-driven cancers.

### Cross-Species Validation of JNK, Toll, Notch, and Hippo Pathways Oncogenic Potentials

With the idea of JNK, NF-κB/Toll, Notch, and Hippo pathways co-operation may be a shared mechanism to drive tumorigenesis, we integrated publicly available single-cell datasets from patients with *KRAS/NRAS-*driven cancers (CRC, LUAD, PAAD, and melanoma) and obtained 174,385 high-quality cells (Fig. 7A) ^76–81^. Malignant epithelial cells were isolated based on lineage markers and further stratified into subpopulations (Figs. S13A-D). We scored the corresponding pathway activity and observed heterogenous regulation of RAS, MAPK, JNK, Toll-like receptor (TLR), Notch, and Hippo signaling across tumors, with subclusters showing co-activation of multi signaling (Figs. S14A-D, Supplementary Table_S3). This led us to further verify the existence of co-activation of these signaling pathways in *RAS*-activated cells (mimicking cells carrying *KRAS/NRAS* oncogenic mutations). By selecting the subclusters with top-ranking RAS signaling for correlation analysis (Fig. S15A), we found that, within *RAS*-activated cells, RAS signaling positively correlated with MAPK, JNK, TLR, and Notch activation and Hippo inactivation in all cancers, except for TLR in CRC and Notch in PAAD (Fig. S15B). This indicated a conserved co-activation pattern mirroring *Drosophila Ras^V12^; fry^⁻/⁻^* tumors.

**Figure 7.**
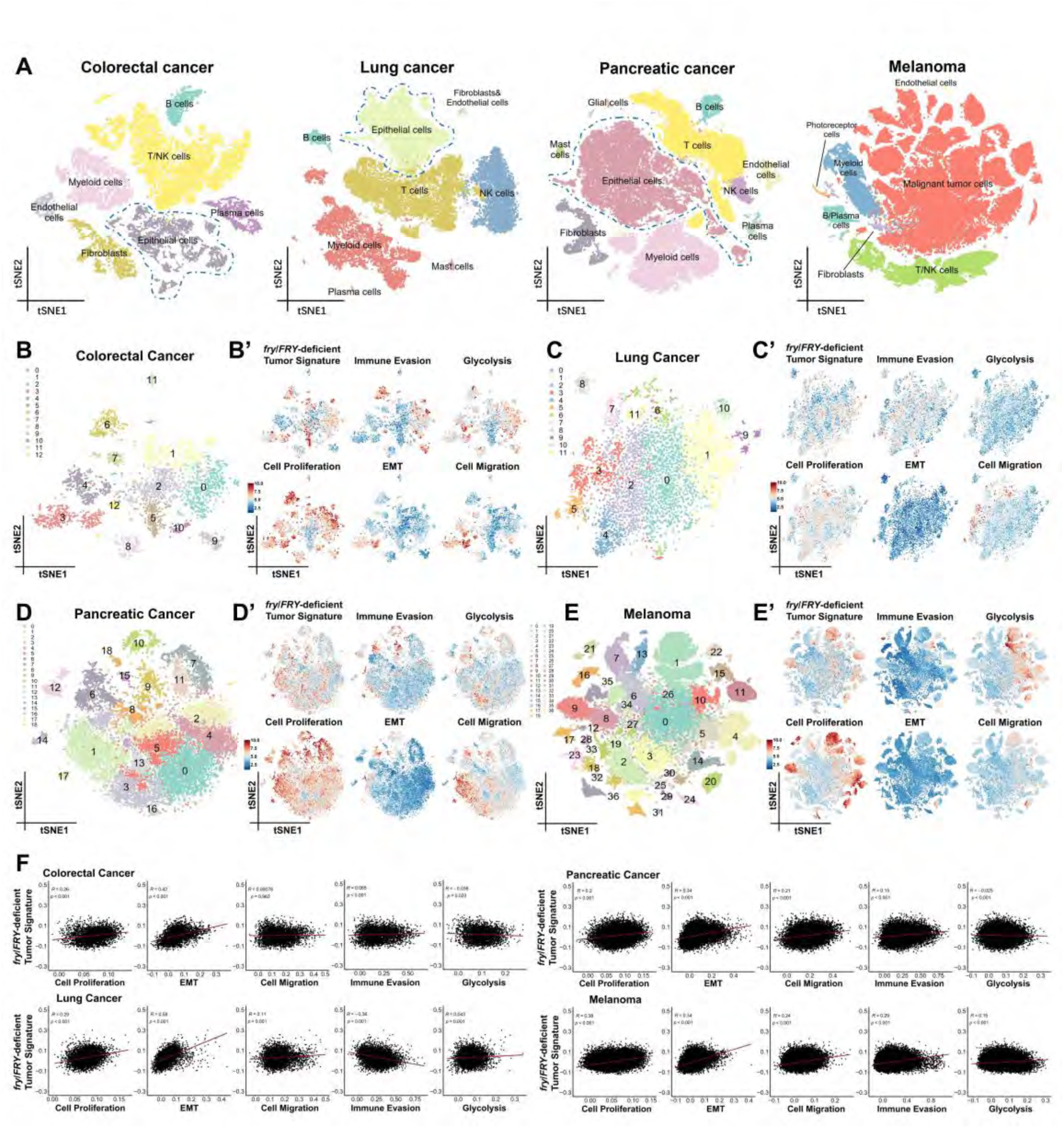
**Cross-species validation of evolutionarily conserved oncogenic mechanisms from *Drosophila* to humans.** (A) Single-cell data analysis for human *KRAS*/*NRAS*-driven cancers. A total of 174,385 high-quality cells were maintained, with 18,727 from colorectal cancer (CRC carrying *KRAS* oncogenic mutations), 25,059 from lung cancer (LUAD), 45,747 from pancreatic cancer (PAAD carrying *KRAS* oncogenic mutations), and 84,825 from melanoma, respectively. The corresponding cell type was annotated. (B-E’) The tSNE projections of re-clustered malignant tumor cells or tumor epithelial cells derived from *KRAS*/*NRAS*-driven cancers, CRC (B), LUAD (C), PAAD (D), and melanoma (E). The tSNE representations depict the scoring of tumor malignancies in re-clustered tumor epithelial cells, CRC (B’), LUAD (C’), PAAD (D’), and melanoma (E’). The evaluation of tumor malignancies encompasses *fry*/*FRY*-deficient tumor signature, cell proliferation, EMT, cell migration, immune evasion, and glycolysis. The scoring values were scaled from 0 to 10. (F) The correlation scatter plots illustrate the relationships between *fry*/*FRY*-deficient tumor signature scores and tumor malignancies scores across *KRAS*/*NRAS*-driven cancers, CRC, LUAD, PAAD, and melanoma. These correlations were determined using Spearman’s correlation analysis, with coefficients R and *P-*values reported, where *P* < 0.05 indicates statistical significance.

To illustrate the relationships between co-activation patterns and tumor malignancies, we employed ‘*fry*/*FRY*-deficient tumor signature’, summarized from JNK, TLR, and Notch activation and Hippo inactivation signatures, alongside signatures reflecting cell proliferation, EMT, cell migration, immune evasion, and glycolysis to systematically evaluate the tumor malignancies (Figs. 7B-E’, Supplementary Table_S3). The *fry*/*FRY*-deficient tumor signature positively correlated with RAS signaling across cancers (R >0.6) (Fig. S16A), suggesting its potential as a positive marker in *RAS*-hypermutated cancers. Overall, *fry*/*FRY*-deficient tumor signature positively correlated with cell proliferation, EMT, and cell migration (except for CRC) across four cancers. The correlations with immune evasion were negative in LUAD, while positive in PAAD and Melanoma. Although the correlation was extremely weak, this signature was significantly associated with glycolysis in all cancers (Fig. 7F). Similar results were observed in *RAS*-activated cells only, where *fry*/*FRY*-deficient tumor signature was also positively correlated with cell proliferation (except for CRC) and EMT, yet weakly and negatively correlated with glycolysis in all cancers (Fig. S16 B-E). Additionally, the correlations with immune evasion were negative in CRC and LUAD while positive in PAAD and Melanoma, consistent with overall findings, suggesting *FRY* may play a dual role in regulating interactions between tumor cells and immune cells (Fig. S16 B-E). Collectively, these results reveal an evolutionarily conserved transcriptomic signature in *RAS*-driven tumors, where JNK, TLR, and Notch activation synergize with Hippo inactivation to fuel malignancy, highlighting their universal relevance in tumor progression.

To further elucidate the tumor-suppressive role of *fry/FRY* in human cancers, we divided six *KRAS*/*NRAS*-driven tumor samples into *FRY*-high and *FRY*-low groups based on the median expression levels of *FRY* for tumor purity and immune infiltration analysis. The analysis of tumor purity using ESTIMATE^82^ revealed that, stromal cell scores, immune cell scores, and Estimate scores were significantly higher in five cancers (COADREAD, LUADLUSC, SKCM, STAD, and UCEC) in *FRY*-high group compared to *FRY*-low group (except for immune cell score in UCEC) (Fig. S17A). This suggests that the high *FRY* expression may induce increased immune cell infiltration, elevated stromal cell counts, and decreased tumor purity.

The detailed infiltration of immune cells was investigated via CIBERSORT ^83^, and we found that low *FRY* expression resulted in common features of increased infiltrations of activated mast cells, CD4 T follicular helper (TFH) cells, activated CD4 memory T cells, and CD8 T cells, along with the decreased M2 macrophages and naïve B cells (Fig. S17B), suggesting that downregulating *FRY* may activate mast cells to promote tumor overgrowth, while activate TFH cells to support B cell functions and further promote the CD8 T cell anti-tumor immunity. This hypothesis was supported by a murine LUAD model, where T cell-B cell interactions promote the tumor-specific TFH cells to secrete interleukin (IL)-21 to support the anti-tumoral function of CD8 T cells ^84^. However, the interactions between *FRY*-low tumor cells and various immune cells require further explorations to elucidate.

### Machine Learning Reveals the Clinical Relevance of DRTs Features

To explore the diagnostic value of tumor suppressors in DRTs for human cancers, we conducted ROC diagnostic analyses on human homologous genes (*FRY* and *SCRIB*) with high mutation rates. We found that *FRY* gene exhibited excellent diagnostic performance for LUAD, LUSC, PAAD, and UCEC (AUC > 0.85), while *SCRIB* exhibited excellent diagnostic performance in COAD, READ, and LUSC (AUC > 0.85) (Figs. 8A and 8B). These results highlight the diagnostic value of *Drosophila* tumor models in clinical applications.

**Figure 8.**
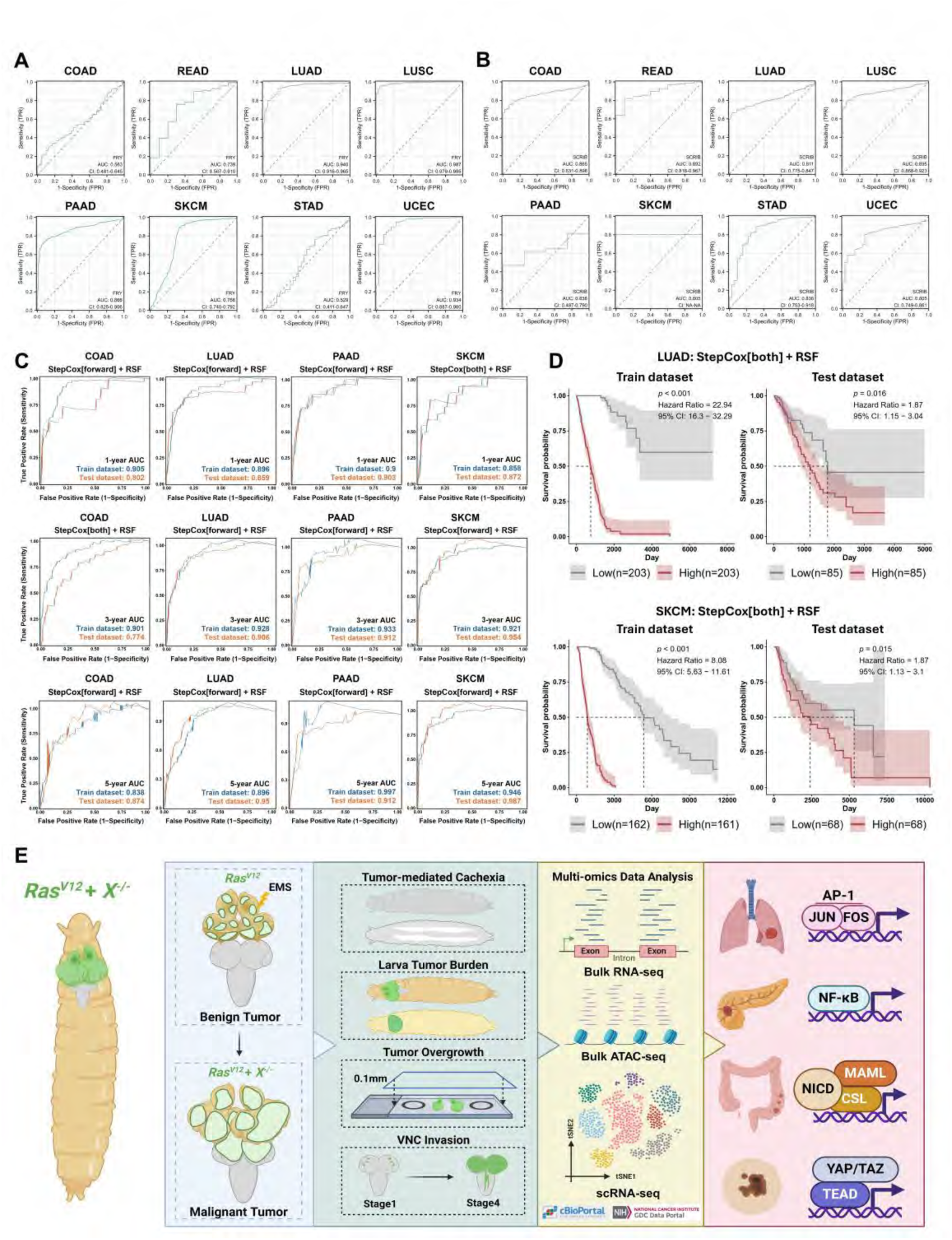
**Machine learning evaluation of predictive value of *Drosophila* tumor model in patient survival across *KRAS*/*NRAS*-driven cancers.** (A) The ROC evaluation of *FRY* diagnostic value across *KRAS*/*NRAS*-driven cancers. The AUC values of each cancer were presented with confidence intervals. The AUC higher than 0.85 was considered excellent performance. (B) The ROC evaluation of *SCRIB* diagnostic value across *KRAS*/*NRAS*-driven cancers. The AUC values of each cancer were presented with confidence intervals. The AUC higher than 0.85 was considered excellent performance. (C) The optimal machine learning model for predicting the survival of 1-year (Top), 3-year (Middle), and 5-year (Bottom) of *KRAS*/*NRAS*-driven cancer patients. The selected machine learning model was denoted under the cancer type. The AUC values of training datasets and test datasets were presented. The AUC higher than 0.85 was considered excellent performance. (D) The applications of StepCox[both] + RSF model in predicting the overall survival of patients with LUAD and SKCM. *P* < 0.05 indicates statistical significance. (E) The proposed model of this study. The model illustrated the utilization of ART-D phenotypic malignancies and dynamic transcriptomic profiles to uncover the oncogenic mechanisms driving tumorigenesis in *Drosophila Ras*-driven tumors. Subsequent integrated multi-omics data analysis revealed an evolutionarily conserved role of oncogenic signaling pathways in human *KRAS*/*NRAS*-driven cancers. The cross-species findings demonstrated strong performance in diagnosis and prediction of human cancers. By integrating *Drosophila* genetics with human oncology, ART-D offers a valuable resource for identifying tumor stage-specific therapeutic targets and advancing the treatment of RAS-driven malignancies.

Furthermore, we seek to exploit the highly conserved nature of signaling pathways to establish optimal prognostic prediction models in various *KRAS*/*NRAS* cancers. To do this, we randomly divided TCGA samples of COAD, LUAD, PAAD, and SKCM into training (70%) and testing sets (30%). Using machine learning methods, we selected Drosophila tumor suppressors human orthologs and key components of signaling pathways as candidate genes (Supplementary Table_S4). After filtering for prognostic features through univariate Cox regression analysis (Supplementary Table_S5), we used 117 combinations of machine learning algorithms from Mime^85^ to develop prognostic prediction models tailored to different human *KRAS*/*NRAS* cancers. By evaluating the performance of each model by calculating the ROC AUC values for 1-year, 3-year, and 5-year survival times in both the training and testing sets (Figs. S18-S21), we elicited the optimal predictive models and found that gene sets based on features of DRTs performed well in predicting 1-year, 3-year, and 5-year survival outcomes for patients with COAD, LUAD, PAAD, and SKCM (AUC > 0.75), with particularly excellent performance for LUAD (AUC > 0.85), PAAD (AUC > 0.9), and SKCM (AUC > 0.85) patients (Fig. 6C).

Notably, the combinations of random forest (RSF) and stepwise Cox (StepCox), StepCox[forward] + RSF and StepCox[both] + RSF, performed well in predicting 1-year, 3-year, and 5-year survival outcomes for COAD, LUAD, PAAD, and SKCM patients. Thereby we intersected the top five machine learning models ranked by 1-year, 3-year, and 5-year survival prediction performance for each cancer to identify models that performed excellently across all three survival time predictions and used these models to predict OS (Figs. S17-S20). Using the prediction model generated by the StepCox[both] + RSF algorithm,

LUAD and SKCM patients were divided into high-risk and low-risk groups. Results showed that the OS of patients in the high-risk group was significantly lower than that of patients in the low-risk group (Fig. 8D), indicating the StepCox[both] + RSF algorithm can effectively predict the survival prognosis of *KRAS*/*NRAS* cancer patients based on features of DRTs.

## Discussion

In this study, we establish the *Drosophila* Atlas of Ras-driven Tumors (ART-D), a relative comprehensive resource that systematically maps the dynamic progression of malignancy across ten genetic contexts. Using ART-D, we clearly define different stages of tumorigenesis: initial stage, promotion stage, and progression stage. Although all DRTs ultimately develop aggressive phenotypes, their progression trajectories vary significantly. Therefore, employing robust linear regression, we classify DRTs into three distinct subtypes: Group A, B, and C. By integrating longitudinal phenotypic profiling with transcriptomic and multi-omics analyses, we delineate a cross-species unified framework for tumor progression. We identify coordinated dysregulation of key signaling pathways, JNK, Notch, NF-κB/Toll, and Hippo, emerge as central drivers of malignancy, with this dysregulation showing striking parallels to human cancers harboring frequent *RAS* mutations. Using machine learning to construct optimal prognostic prediction models tailored for human *KRAS*/*NRAS* cancers, we demonstrate that these evolutionarily conserved oncogenic mechanisms hold significant potential for clinical applications. (Fig. 8E).

Tumor progression involves dynamic accumulation of genetic and epigenetic alterations that collectively drive phenotypic evolution ^86–88^. The ART-D model advances cancer research by enabling real-time dissection of these dynamics in a genetically tractable system. Unlike clinical samples that offer static snapshots, ART-D captures three distinct progression stages. The formation rates of features at each stage vary, so we innovatively applied a robust regression model to classify their phenotypic characteristics. This approach, combined with their dynamic transcriptomic features, enabled us to explore potential molecular mechanisms that promote tumor malignancy progression, an approach rarely used in Drosophila tumor research, bridging the gap between genetic lesions and emergent phenotypes. This resource provides critical insights into temporal dynamics of tumor progression, positioning *Drosophila* as a powerful platform for studying tumor progression mechanisms.

Importantly, our study uncovers profound conservation of tumorigenic mechanisms between *Drosophila* and humans. Cross-species mapping of tumor suppressors identifies *SCRIB*, *LLGL1/LLGL2*, and *FRY* as key genes frequently dysregulated in *KRAS/NRAS*-mutant human cancers, where their loss correlates with poor patient survival. For example, loss of *SCRIB* synergizing *KRAS^G12D^* induces aggressive lung tumors by activating MAPK signaling ^89^, consistent with *Drosophila* models ^27^. Besides, *SCRIB* deficiency drives ductal hyperplasia in mammary models and cooperates with *KRAS^G12D^*and *P53* loss to accelerate pancreatic adenocarcinoma metastasis ^90^. Notably, *SCRIB* mislocalization also underlies adaptive resistance to *KRAS^G12C^* inhibitors, underscoring its clinical relevance ^91^. While earlier work highlighted pairwise pathway interactions (e.g., JNK-Hippo, Hippo-Notch) ^35–38, 92, 93^, we identify a four-pathway nexus in *Ras^V12^; fry^−/−^* tumors: synergistic activation of JNK, Notch, and NF-κB/Toll alongside Hippo inactivation. Strikingly, single-cell transcriptomics and machine learning investigation of *RAS*-mutant human cancers (pancreatic, colorectal, lung, melanoma) reveal analogous co-activation of these pathways, suggesting an evolutionarily conserved network driving malignancy while revealing pathway interdependencies that may represent therapeutic vulnerabilities.

Although our study provides systematic and comprehensive insights into *Ras*-driven tumorigenesis, several limitations remain. First, while the *Drosophila* model excels in genetic resolution, it lacks key components of the mammalian tumor microenvironment, such as adaptive immunity and interactions with stromal cells. Incorporating immune responses and co-culture systems into future studies may enhance translational relevance. Second, this study focused on oncogenic *Ras* mutations and their cooperation with specific combinations of tumor suppressor gene alterations to drive malignant phenotypes. This provides a foundation for further investigation into broader oncogenic and tumor suppressive pathways to uncover more general mechanisms of tumorigenesis. Additionally, although we identified evolutionarily conserved synergistic interactions among pro-tumorigenic signaling pathways from *Drosophila* to humans, the precise molecular mechanisms regulating the JNK, Notch, NF-κB, and Hippo pathways remain unclear. Finally, functional validation in human cancer models, particularly patient-derived organoids and xenograft models, is still lacking, which is crucial for assessing the therapeutic potential of targeting these pathways.

In summary, through systematic investigation of *Ras*-driven tumors in *Drosophila*, we have developed a comprehensive platform, ART-D, that characterizes the progressive phenotypes and dynamic transcriptomic features of *Ras*-driven malignancies. Our study reveals conserved cooperative mechanisms among JNK, Notch, NF-κB, and Hippo signaling pathways from *Drosophila Ras* tumors to human *KRAS*/*NRAS* cancers, establishing a systematic framework for cross-species translational cancer research. By integrating *Drosophila* genetics with human oncology, ART-D represents a valuable resource for identifying stage-specific therapeutic targets and advancing the treatment of *RAS*-driven cancers.

## Materials and Methods

### Reagents and tools table

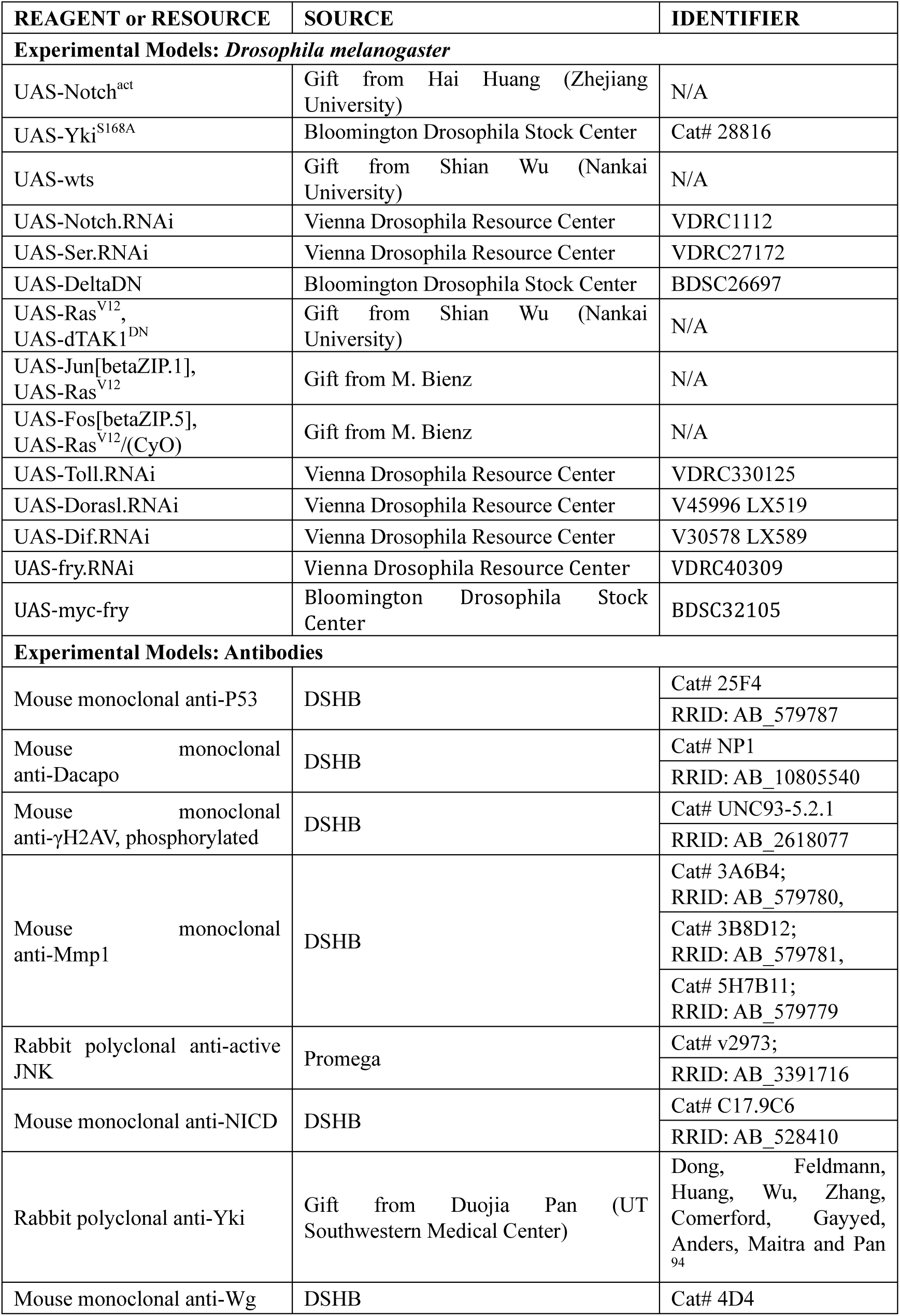
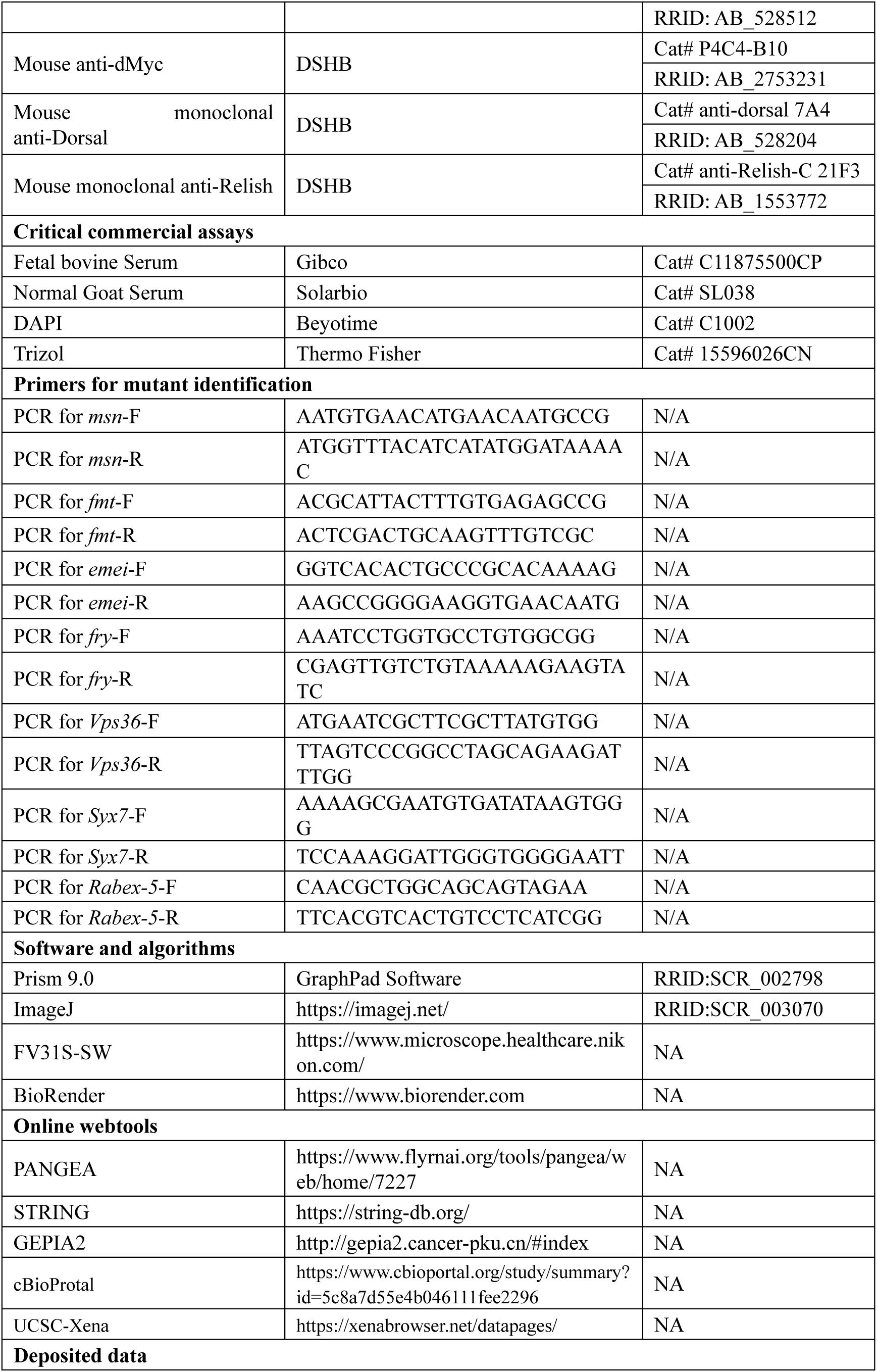
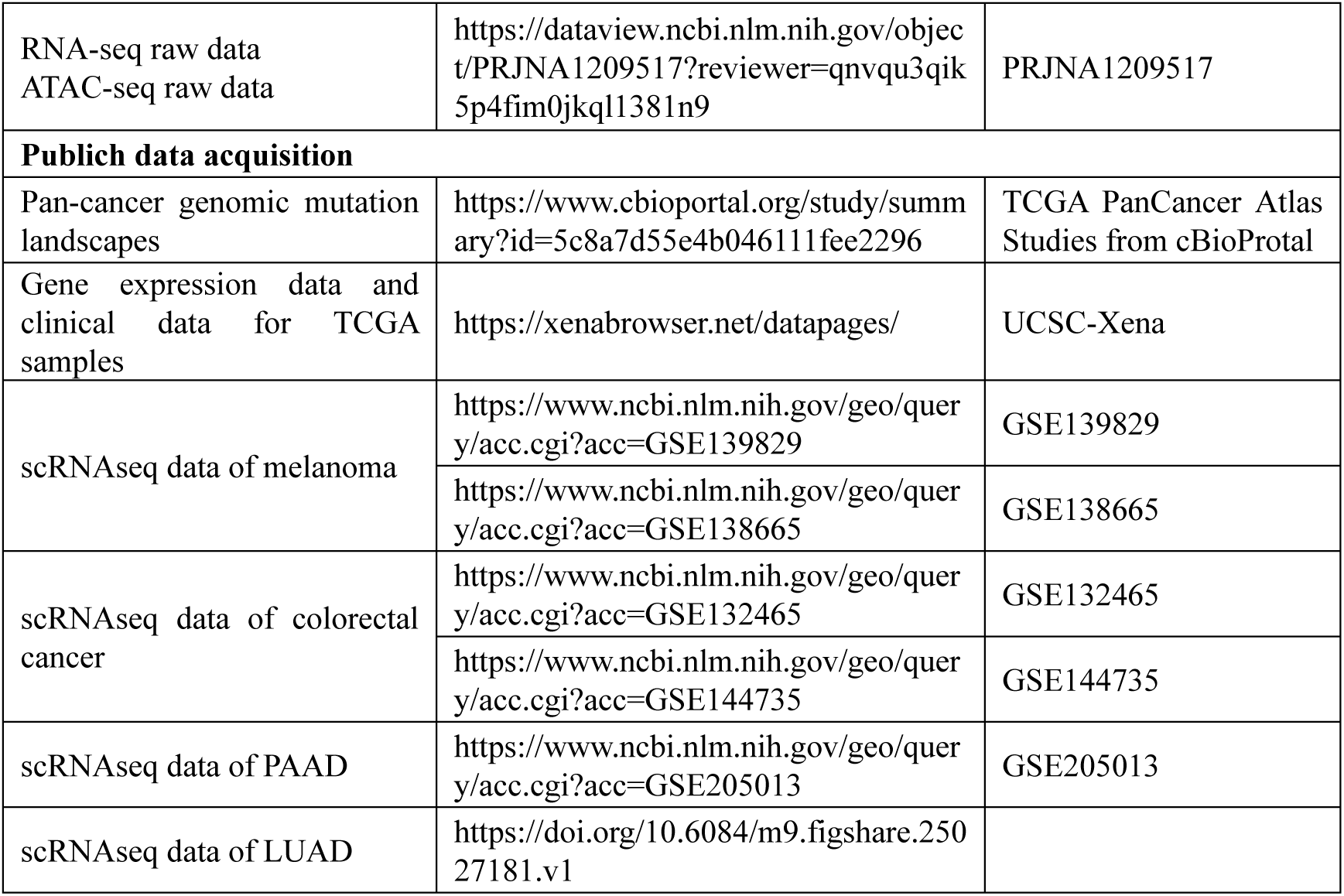

### Fly husbandry and genetics

*Drosophila* stocks and crosses were maintained on standard food at 25 °C under standard laboratory conditions (25 °C, 12:12 h light: dark). Standard cornmeal-yeast food containing 50 g corn flour, 24.5 g dry yeast, 9 g agar, 7.25 g white sugar, 30 g brown sugar, 4.4 ml propionic acid, 12.5 ml ethanol and 1.25 g nipagin per liter. A comprehensive list of the utilized *drosophila* strains is provided in the reagents and tools table.

### EMS screen and mutant identification

EMS Screen details were described previously^35^. A benign tumor model carrying the *Ras^V12^* was established in the eye-antennal imaginal discs of *Drosophila* using the MARCM system. Single-gene mutations were induced in parent *Drosophila* (*UAS-Ras^V12^*; *79E*) through EMS treatment. After crossing with *79E* tester flies, homozygous mutant clones were generated specifically in the *Ras^V12^* mutant cells. Novel tumor suppressor genes were identified based on the size of GFP-labeled regions and the tumor burden observed in the eye-antennal imaginal discs of the *Drosophila* larvae. By crossing *Drosophila* deficiency lines with mutant fly strains of candidate tumor suppressor genes, when homozygous lethality occurs, mutant alleles of specific genes are used for precise localization. For each mutant, *Drosophila* lines were examined by Sanger sequencing. Genomic DNA was rapidly extracted using squish buffer (10 mM Tris-Cl, pH 8.2, 1 mM EDTA, 25 mM NaCl, 200 µg/ml Proteinase K) and then added to the PCR reaction to amplify the target gene segment. The mutation sites were subsequently confirmed by Sanger sequencing. The primers used for PCR were listed in the reagents and tools table.

### Phenotypic malignancies acquisition

The four representative phenotypic characteristics of normal tissue, *Ras* benign tumors, and various malignant tumors were harvested at different time points (5.5d ± 6h, 9.5d ± 6h, and 13.5d ± 6h AEL) as indicated. Whole larvae were photographed using a fluorescent stereomicroscope (Olympus, SZX16) with GFP fluorescence or bright field to quantify larval tumor burden and tumor-mediated cachexia. The ratio of GFP regions to the whole larva highlights the tumor growth burden on the host, while the overall transparency of the larvae reflects the degradation of fat bodies and muscles due to cachexia. Paired larvae were dissected in cold PBS, and the eye-antennal discs or tumors were separated from the VNC and brain tissues as much as possible. The eye-antennal discs or tumors were placed in a chamber with a 0.1 mm height to assess overall tissue volume or GFP region volume, quantifying tumor overgrowth. The relative occupation rate of the VNC, as a neighboring tissue, reflects the invasion ability of tumor cells. Quantifications of these images were performed using ImageJ software.

### Larval pupariation rate quantification

Roughly ten males were crossed with twenty females per vial, and the vials were changed daily. The number of pupae per vial was counted at the indicated time points. The mean number of pupae was calculated for each genotype by averaging the results from three independent replicate vials.

### Immunofluorescent staining and imaging

The imaginal discs were dissected from the third-instar larvae on the indicated days AEL. Dissections were performed in cold PBS, and the discs or tumors were fixed with 4% paraformaldehyde for 15 minutes at room temperature with gentle shaking. Subsequently, the samples were washed three times with PBS containing 0.1% Triton X-100 (PBST) for 5 minutes each. After fixation and washing, samples were blocked in 10% goat serum in PBST for 30 minutes, followed by incubation with primary antibody (1:200) at 4°C overnight. The primary antibodies used in this study are listed in the reagents and tools table. Samples were then washed three times with PBST for 10 minutes each at room temperature, followed by incubation with secondary antibody and DAPI (1:500 in PBST) for 2 hours at room temperature. This was followed by three additional washes with PBST. The imaging of all samples was conducted using an Olympus FV3000-BX63 confocal microscope (inverted) equipped with 20x or 40x objectives. Images were processed using Olympus FV31S-SW to add scale bars, ImageJ for quantification, and Photoshop CS5 (Adobe) for image merging and resizing.

### Regression analysis and PCA of *Drosophila Ras*-driven tumors

A robust linear regression model (“rlm” method) from MASS (v7.3-65) was applied to construct regression models for the numerical variables of tumor burden, tumor overgrowth, and tumor-induced cachexia in *Drosophila Ras*-driven malignant tumor progression. Scatter plots were generated using ggplot2 (v3.4.4), and regression equations were added using ggpmisc (v0.6.1).

Based on robust linear regression equations of different tumor types, the slopes, intercepts, and adjusted R^2^ values from both individual tumor types and the overall *Ras*-driven tumor robust regression model were extracted for PCA analysis. The ten *Drosophila* tumor models were projected onto the two-dimensional interface formed by PC1 and PC2, and K-means clustering was applied to classify DRTs into distinct subtypes. Additionally, the contributions of PC1 and PC2 to numeric variables reflecting tumor malignancy progression, including larval tumor burden, tumor overgrowth, and tumor-induced cachexia, were visualized using heatmaps generated with pheatmap (v1.0.12) and RColorBrewer (v1.1-3).

### Total RNA extraction

For control groups, a total of 200 eye-antennal discs (wildtype group, 3 replicates) or benign tumors (Ras group, 3 replicates) were harvested at the late stage of third instar larvae for total RNA extraction by using Trizol (Invitrogen, Carlsbad, CA) according to manual instruction. For case groups, a total of 100 eye-antennal malignant tumors from *Ras^V12^//X^-/-^*(case group, 3 replicates) were harvested at three different time points (5.5d, 9.5d, and 13.5d AEL) for total RNA extraction with the same procedures. The RNA samples were subjected to Nanodrop 2000 (Thermo Fisher Scientific) and 2100 bioanalyzer (Aglient) for quality control. Samples with concentrations higher than 30 ng/µl, OD260/OD230 ratios between 2.0 and 2.5, and RNA integrity number (RIN) values greater than 8.0 were considered as qualified samples for further library preparation.

### Library preparation and RNA-seq

For each sample, a total amount of 1-3μg RNA was used as the input for library preparation by strictly following the standard protocol of VAHTS Universal V6 RNA-seq Library Prep Kit for Illumina^®^ (NR604-01/02). Briefly, mRNA was purified by using poly-T oligo-attached magnetic beads from total RNA. The short fragments of mRNA were obtained by adding the fragmentation buffer. After the first strand of cDNA was synthesized by using random hexamer primer and RNase H, the second strand synthesis was performed subsequently by using buffer, dNTPs, DNA polymerase I, and RNase H. And then, the double-stranded cDNA was purified by using QiaQuick PCR kit or AMPure P beads. The purified products of each sample were repaired at the end, added tail, and connected to the sequencing connector, then the appropriate fragment size was selected, and the final cDNA library was obtained by PCR amplification for further sequencing performed on Illumina NovaSeq 6000 platform with NovaSeq 6000 S4 Reagent kit V1.5.

### RNA-seq data processing and functional annotation

All eye-antennal disc/tumor transcriptomic data were processed and analyzed as follows. FastQC (v0.11.8) (https://github.com/s-andrews/FastQC/releases/tag/v0.11.8) software was applied for data quality control and reads statistics. The high-quality reads (Q30 > 90%) were mapped to the Ensemble ^95^ *Drosophila melanogaster* reference genome (Drosophila_melanogaster.BDGP6.32.108) by using Hisat2 ^96^, followed by the gene feature counting with default settings by using HTSeq ^97^. The DEGs identification and over-representation analysis (ORA) were conducted in R (v4.2.0) after removing batch effect using the *ComBat*_seq function of sva (v3.46.0) package ^98^. The gene count matrix after batch effect removal was provided (Supplementary Table_S1). Genes meeting the criteria of |Fold-Change| >1.5 and FDR <0.05 were identified as differentially expressed genes (DEGs) with the “RLE” method of edgeR (v3.40.2) ^99^. The gene id transformation was performed by using biomaRt ^100^. The ORA of DEGs were performed against terms in various databases, including FlyBase signaling pathway (https://flybase.org), Kyoto Encyclopedia of Genes and Genomes (KEGG) database ^101, 102^, Gene Ontology (GO) consortium ^103^, and Drosophila RNAi Screening Center (DRSC) PathON database (https://www.flyrnai.org/tools/pathon/web/) by using PANGEA webtools ^104^ or clusterProfiler package ^105^. Data visualization was relied on limma (v3.54.2), pheatmap (v1.0.12), RColorBrewer (v1.1-3), and ggplot2 (v3.4.4) packages.

### Linking phenotypic data with transcriptomic data

WGCNA (v1.73) was applied to identify modules of highly correlated genes and associate the modules with sample phenotypic traits ^45^. After pre-process of sample data to remove the outliers, the optimal soft threshold was chosen to convert the correlation matrix into an adjacency matrix. Modules were generated by using the *blockwiseModules* function. The cluster dendrograms were recorded and plotted to show the correlations of different modules. The Pearson correlation analysis was performed to screen the modules that were significantly associated with parameters of four representative phenotypic characteristics or genotypes. A *P* value less than 0.05 was considered significant. The modules of interest were selected for further analysis.

### Time-series analysis of transcriptomic data

The gene count matrix was transformed into Log2(cpm+1) for input. Two packages, Mfuzz (v2.58.0) and ClusterGVis (v0.1.2), were utilized to illustrate the gene dynamic change patterns across five pseudotime points, simulating the process of tumor progression from normal to malignant. For Mfuzz ^46^, the m value was estimated using the *mestimate* function, and genes with memberships below 0.2 were excluded. For ClusterGVis (https://github.com/junjunlab/ClusterGVis), K-means clustering was applied to generate 10 clusters, and the dynamic gene expression patterns were visualized using heatmaps with key genes highlighted.

### Protein and protein interaction network

Genes of interest were used as input to construct PPI networks using the STRING database ^47^, which were subsequently visualized in Cytoscape software ^106^. The selected genes were evaluated using 11 algorithms within the cytoHubba plugin, and the Bottleneck algorithm was chosen to identify hub genes ^107^. The top 25 hub genes were then selected.

### Transcription factors enrichment analysis

The enrichment of transcription factors was conducted by using RcisTarget (v1.26.0) ^108^. The motifAnnotations_dmel was adopted for the annotations of motifs to TFs. The rankings of motif were set by using *importRankings* function to import the specie corresponding file (dm6_v10_clust.genes_vs_motifs.rankings.feather) downloaded online (https://resources.aertslab.org/cistarget/).

### GSVA and ssGSEA

A total of 15 signaling pathways were selected, including lnR, Hippo, Notch, Toll, JAK_STAT, TNFα_Eiger, FGFR, Hedgehog, Pvr, Wnt, Sevenless, Imd, Torso, BMP, and EGFR signaling pathways. For each signaling pathway, the list of positive regulators, negative regulators, and all members were collectively downloaded from FlyBase (https://flybase.org) ^109^. The GSVA ^48^ and ssGSEA ^49, 50^ were performed locally using the gene lists with GSVA (v1.46.0). The average values of three replicates from each sample were subjected to Pearson correlation analysis with the average values of the basic phenotypes of tumor progression. A *P* value less than 0.05 was considered statistically significant.

### Library preparation and ATAC-seq

Either disc or tumor samples were digested into a single-cell suspension using the sCelLive^TM^ Tissue Dissociation Mix (Singleron). Cell viability and concentration were assessed using Trypan Blue staining (Cat#C0040, Solarbio) and Acridine Orange/Propidium Iodide (AO/PI) staining (#RE010212, Countstar). For each sample, a total of 2 × 10^5^ cells were used for ATAC-seq library preparation. The library preparation was conducted using the Hyperactive ATAC-Seq Library Prep Kit for Illumina (Vazyme, TD711) according to the manufacturer’s guidelines. Cells were centrifuged at 500 × *g* at 4 °C for 5 min, resuspended, and washed twice with 50 μl of TW Buffer. The cells were then resuspended in 50 μl of cold lysis buffer and incubated on ice for 5 min to isolate the nuclei. Following a 10 min centrifugation at 500 × *g* at 4 °C, the nuclei were incubated in a 50 μl fragmentation reaction mix at 37 °C for 30 min. The desired DNA was extracted using 100 μl of ATAC DNA Extract Beads, followed by two washes with 200 μl of fresh 80% ethanol and a final elution with 26 μl of nuclease-free ddH_2_O. Library amplification was performed by adding 60 μl of PCR mix to 20 μl of DNA product, following the program: 72 °C for 3 min; 95 °C for 3 min; 12 cycles of 98 °C for 10 s, 60 °C for 5 s, and 72 °C for 1 min; and a hold at 12 °C. The amplified ATAC-Seq library was further purified using ATAC DNA Clean Beads, followed by two washes with 200 μl of fresh 80% ethanol and a final elution with 22 μl of nuclease-free ddH2O. The final ATAC-seq libraries were sequenced using the Illumina NovaSeq 6000 platform.

### ATAC-seq data processing and analysis

The analytic procedures of ATAC-seq data were referred to recommended pipelines ^110^. The sequencing data obtained from the Illumina platform was processed by using FastQC (v0.11.8) software for quality control. Adapters were removed and reads were trimmed by using Cutadapt (v1.18) ^111^. The clean reads were first mapped to the Ensemble ^95^ *Escherichia coli* (E. coli) reference genome (GCF_000005845.2_ASM584v2_genomic) by using Bowtie2 (v2.4.2) ^112^, the unrecognized reads were subsequently mapped to *Drosophila melanogaster* reference genome (Drosophila_melanogaster.BDGP6.32.108) with the parameters “--end-to-end --very-sensitive --no-mixed --no-discordant --phred33 -I 0 -X 2000 --no-unal”. After using SAMtools (v1.11) ^113^ to transform SAM files into BAM files, Picard (v2.25.1) (https://broadinstitute.github.io/picard/) was applied to remove duplicates, and the reads were selectively maintained when meeting “mapping_quality >= 20”. The final BAM files of same genotype among the replicates were merged into one for peak calling, which is performed by using Model-based Analysis of ChIP-seq (MACS2, v2.2.6) ^114^ with settings “-g dm -f BAMPE --nomodel --shift -100 --extsize 200”. Peaks were annotated by using ChIPseeker ^115^ package in R (v4.2.0), and the BAM files were transformed into bigwig files for signal enrichment visualization by using the *bamCoverage* function of BEDTools (v2.30.0) ^116^ with settings “--binSize 20 --normalizeUsing RPKM”. The average normalized signal values of peaks in *drosophila* genome region were calculated and visualized by using DeepTools (v3.5.1) ^117^ and IGV ^118^, respectively. The binding motifs were identified by using the *findMotifsGenome* or *findMotifs* functions of Homer (v4.11) ^119^ regarding to conditions.

### Pan-cancer evaluation of *Drosophila* tumor suppressor ortholog mutation

The *Drosophila* tumor suppressor orthologs were identified using various databases ^62–75^. As a result, a total of 18 human orthologs were identified corresponding to ten *Drosophila* tumor suppressors. The mutation rates of *RAS* family genes (*KRAS*, *NRAS*, and *HRAS*) and *Drosophila* tumor suppressor orthologs were analyzed by using the pan-cancer genomic data obtained from cBioPortal ^60^. Samples were divided into two groups based on the target gene mutations, then prognostic impact was evaluated by performing Kaplan-Meier analyses with Log-rank test.

### Pan-cancer expression and prognosis of *Drosophila* tumor suppressor orthologs

The pan-cancer expression levels of 18 *Drosophila* tumor suppressor orthologs were investigated using a Log_2_(TPM+1) transformed gene expression matrix, downloaded from UCSC-Xena ^61^, encompassing TCGA and GTEx samples. The prognostic value of these human orthologs was assessed locally through Kaplan-Meier analyses with Log-rank tests, and via GEPIA2 online ^120^, employing appropriate cut-offs to delineate high and low group samples.

### Immune infiltration analysis

The immune infiltration of tumor samples from TCGA *KRAS*/*NRAS*-driven cancers were evaluated by using ESTIMATE^82^ and CIBERSORT^83^, respectively. Tumor samples obtained from TCGA were divided into two groups based on the medial expression level of *fry* ortholog gene *FRY*. The tumor purity was evaluated by using ESTIMATE. The relative infiltration level of various immune cell subpopulations were calculated by using CIBERSORT ^83^. The relationships between *FRY* expression and immune infiltrations in *KRAS*/*NRAS*-driven caners were calculated via Wilcoxon rank sum test.

### scRNA-seq data analysis

Six scRNA-seq datasets encompassing four cancer types: pancreatic cancer (PAAD), melanoma (SKCM), colorectal cancer (CRC), and lung adenocarcinoma (LUAD) were collected ^76–81^. Each dataset included tumor cells, stromal cells, and immune cells. The scRNA-seq datasets were processed and analyzed using the Seurat package (v4.0.0) in R (v4.2.0). The high-quality cells were defined as follows: for CRC and LUAD, cells satisfied the criteria “1500 < *nCount_RNA* < 40000; 200 < *nFeature_RNA* < 6000; the proportion of mitochondrial genes < 15%” were maintained; for SKCM, cells satisfied the criteria “200 < *nFeature_RNA* < 8000; the proportion of mitochondrial genes < 10%” were maintained; for PAAD, cells satisfied the criteria “500 < *nFeature_RNA*; 1500 < *nCount_RNA*; the proportion of mitochondrial genes < 15%” were maintained. The scRNA-seq data from different patients or different datasets were integrated by using canonical correlation analysis (CCA) with the *FindIntegrationAnchors* and *IntegrateData* functions ^121^. *NormalizeData* and *ScaleData* functions were used to normalize and scale gene counts, *FindVariableFeatures* function was applied to select 2500 top variable genes for PCA, and further dimensionality reduction using tSNE with *RunTSNE* function ^122^. Primary cell clustering was performed using the Louvain algorithm with an appropriate resolution. The resulting Louvain clusters were visualized in a two-dimensional tSNE plot, and each cluster was annotated to known biological cell types using canonical marker genes as indicated. The cell types of interest were extracted by using *subset* function, followed by re-performing dimensionality reduction and cell clustering. The visualization of scRNA-seq data was dependent on dplyr (v1.1.4), ggsci (v3.0.0), cowplot (v1.1.1), gridExtra (v2.3), ggpubr (v0.6.0), reshape (0.8.9), tidyverse (2.0.0), RColorBrewer (v1.1-3), and ggplot2 (v3.4.4) packages.

### Scoring and correlation analysis of signaling pathways and tumor malignancies

The scoring of different signaling pathways and tumor malignancies was conducted by using the *AddModuleScore* function with the gene list provided (Supplementary Table_S3). The *fry*/*FRY*-deficient tumor signature encompassed genes from JNK, TLR, Notch activation and Hippo inactivation. The signatures reflecting tumor malignancies were mainly constructed by gene sets recorded in the MSigDB ^123^ and were integrated with gene sets reported in literatures, including cell proliferation (GO_0008283), cell migration (WU_CELL_MIGRATION ^124^), EMT(LIANG_EMT ^125^, KOHN_EMT_EPITHELIAL ^126^, and HALLMARK_EMT), immune evasion (GOURMET_IMMUNE_EVASION ^127^), and glycolysis (HALLMARK_GLYCOLYSIS). The *unique* function was applied to filter the replicated genes. The correlations between two signature gene sets were assessed by using Spearman’s correlation analysis, with coefficients R and *P* values reported, where *P* < 0.05 indicates statistical significance.

### The diagnostic value of *Drosophila* tumor suppressor orthologs

The UCSC-Xena TCGA samples from COAD, READ, LUAD, LUSC, PAAD, SKCM, STAD, and UCEC patients were utilized to evaluate the diagnostic value of Drosophila tumor suppressor orthologs. The gene expression matrix, transformed using log2(TPM+1), was employed, and the diagnostic accuracy of these orthologs was assessed using the pROC (v1.18.0) package to construct ROC curves^128^.

### Optimal prognostic models developed via 117 machine learning combinations

The TCGA samples of COAD, LUAD, PAAD, and SKCM were divided into training dataset (70%) and test datasets (30%) by using the *createDataPartition* function. The candidate gene signature gene list comprises members from JAK-STAT, MAPK, RAS, Hippo, Notch, and TLR signaling pathways (Supplementary Table_S3). The development of prognostic models via machine learning was based on Mime package^85^. The univariate Cox regression analysis was conducted to identify prognostic features as input variables, in which *P* values less than 0.05 considered statistically significant (Supplementary Table_S4). These features were then subjected to a framework comprising 117 machine learning combinations generated from ten classical machine learning algorithms: random forest (RSF), elastic network (Enet), stepwise Cox (StepCox), CoxBoost, partial least squares regression for Cox (plsRcox), supervised principal components (superpc), generalized boosted regression models (GBM), survival support vector machine (survivalsvm), Ridge, and least absolute shrinkage and selection operator (Lasso). The 1-year, 3-year, and 5-year AUC values of ROC were evaluated in both training and test datasets. The top five predictive models based on 1-year, 3-year, and 5-year AUC rankings were combined to identify the optimal model for predicting overall survival in patients with COAD, LUAD, PAAD, and SKCM, respectively. For all ROC analyses, an AUC greater than 0.85 was considered to indicate excellent performance.

### Quantification and statistical analyses

ImageJ (https://imagej.net/ij/) was used to quantify the area of the entire disc, the GFP region, larval transparency, and other parameters. For each group, at least 25 randomly selected samples were used for quantification. Statistical analyses were performed using GraphPad Prism software (version 9.0, GraphPad Software, San Diego, CA). Results are presented as the mean ± SEM or mean ± SD. Comparisons between two groups with a normal distribution and homogeneity of variance were conducted using an unpaired Student’s *t* test; otherwise, the Mann-Whitney *U* test was used. For experiments involving three or more groups, data with a Gaussian distribution of residuals were analyzed using ordinary one-way ANOVA; otherwise, the nonparametric Kruskal-Wallis test was employed. The categorical variables, like larval pupariation rate, were assessed by using the Fisher’s exact test or Chi-square test. The correlation analysis was performed by using Pearson or Spearman’s correlation analysis. And the overall survival was plotted and compared using the Kaplan-Meier method, the patients with different cancers were separated into 2 groups based the gene expression as indicated. The hazard ratios accompanying 95% confidence intervals (CI) in the frost plots were calculated by using Cox proportional hazards model. A *P* value less than 0.05 was considered significant, \**P* < 0.05, \*\**P* < 0.01, \*\*\**P* < 0.001, \*\*\*\**P* < 0.0001.

## Data and code availability

Our bulk RNA-seq and ATAC-seq data for this study have been deposited in the National Center for Biotechnology Informationn’s Sequence Read Archive (PRJNA1209517). Data for TCGA PanCancer Atlas Studies were collected from cBioPortal ^60^. Gene expression data and clinical data for TCGA and GTEx samples were obtained from UCSC-Xena (https://xenabrowser.net/datapages/) ^61^. The scRNA-seq data are available from the Gene Expression Omnibus (GEO) database under accession numbers GSE139829 ^76^, GSE138665 ^78^, GSE132465 ^77^, GSE144735 ^77^, and GSE205013 ^79, 80^, or from Figshare with the identifier https://doi.org/10.6084/m9.figshare.25027181.v1 ^81^. The custom code used in this study is available on GitHub (https://github.com/GYF-pixel/GYF-Drosophila-ART-D.git).

## Acknowledgements

We thank Bloomington *Drosophila* Stock Center, Vienna *Drosophila* Resource Center, and DSHB for providing fly stocks and reagents; the General Equipment and Autoclave Service Core Facility, the Microscopy Core Facility, the Genomics Core Facility, and the High-Performance Computing Center of Westlake University for the facility support. We thank J.L. for conducting the initial genetic screen and Mingfeng Li for critical feedback on the manuscript. The schematic cartoons were created with BioRender.com. This work was supported by grants from the National Natural Science Foundation of China (32170824 and 32322027 to XM); HRHI program (1011103360222B1) of Westlake Laboratory of Life Sciences and Biomedicine to XM; “Pioneer” and “Leading Goose” R&D Program of Zhejiang (2024SSYS0034).

## Author contributions

X.M. conceived, designed, and supervised the study. X.M., Y.G., J.Z. and X.W. designed the experiments and analyzed the data. J.L. conducted the initial genetic screen. Y.G. performed fly-driven experiments with the help from J.Z., K.Z., Y.W., M.W., W.X., W.L., and X.W. K.Z. identified the mutant generated from EMS screening. D.L. helped with the collection of public data. Y.G. performed the multi-omics data analyses. Y.G and X.M. wrote the manuscript.

